# Single nuclei transcriptomics reveal the differentiation trajectories of periosteal skeletal/stem progenitor cells in bone regeneration

**DOI:** 10.1101/2023.06.23.546220

**Authors:** Simon Perrin, Maria Ethel, Vincent Bretegnier, Cassandre Goachet, Cécile-Aurore Wotawa, Marine Luka, Fanny Coulpier, Cécile Masson, Mickael Ménager, Céline Colnot

## Abstract

Bone regeneration is mediated by skeletal stem/progenitor cells (SSPCs) that are mainly recruited from the periosteum after bone injury. The composition of the periosteum and the steps of SSPC activation and differentiation remain poorly understood. Here, we generated a single-nuclei atlas of the periosteum at steady-state and of the fracture site during early stages of bone repair (https://fracture-repair-atlas.cells.ucsc.edu). We identified periosteal SSPCs expressing stemness markers (*Pi16* and *Ly6a*/SCA1) and responding to fracture by adopting an injury-induced fibrogenic cell (IIFC) fate, prior to undergoing osteogenesis or chondrogenesis. We identified distinct gene cores associated with IIFCs and their engagement into osteogenesis and chondrogenesis involving Notch, Wnt and the circadian clock signaling respectively. Finally, we show that IIFCs are the main source of paracrine signals in the fracture environment, suggesting a crucial paracrine role of this transient IIFC population during fracture healing. Overall, our study provides a complete temporal topography of the early stages of fracture healing and the dynamic response of periosteal SSPCs to injury, redefining our knowledge of bone regeneration.

**Graphical Abstract:** 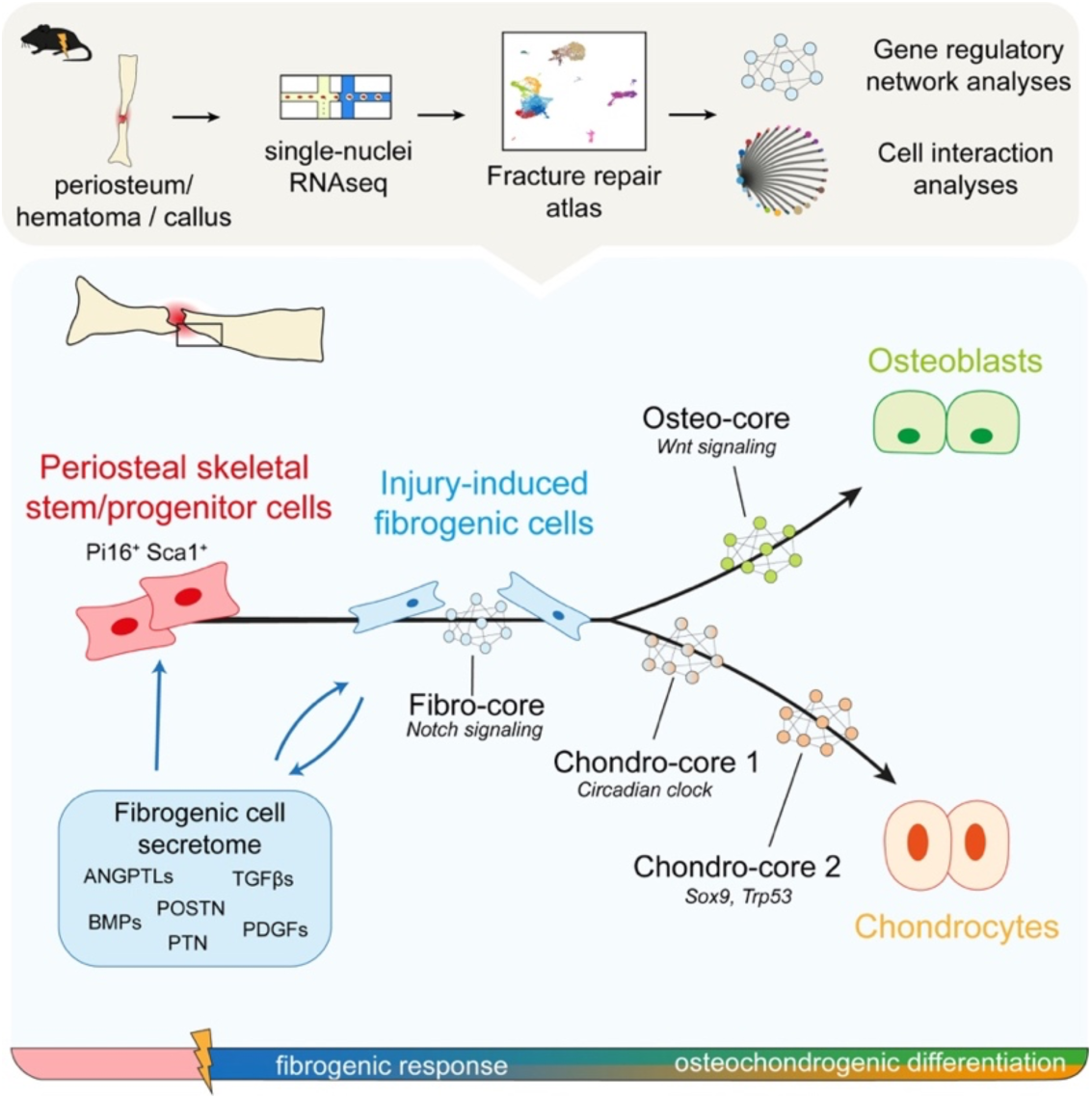

## Introduction

The skeleton provides structural support and protection for internal organs in the vertebrate body. Bones can fracture but regenerate themselves efficiently without scarring. Bone regeneration is mediated by the action of resident skeletal stem/progenitor cells (SSPCs) from periosteum and bone marrow, and SSPCs from skeletal muscle adjacent to the fracture site^1–4^. SSPCs are activated during the inflammatory phase of healing and differentiate into osteoblasts and/or chondrocytes to repair bone via a combination of intramembranous and endochondral ossification. During intramembranous ossification SSPCs differentiate directly into osteoblasts, while during endochondral ossification SSPCs first differentiate into chondrocytes to form an intermediate cartilage template subsequently replaced by bone.

The periosteum, an heterogeneous tissue located on the outer surface of bones, is a crucial source of SSPCs during bone healing^1,2,5–7^. Periosteal SSPCs exhibit a high regenerative potential. They also display both osteogenic and chondrogenic potentials after injury compared to bone marrow SSPCs that are mostly osteogenic and skeletal muscle SSPCs that are mostly chondrogenic^1–4,8^. Periosteal SSPCs (pSSPCs) are still poorly characterized and their response to bone injury remain elusive. Recent advances in single cell/nuclei transcriptomic analyses provided new insights in stem cell population heterogeneity and regeneration processes in many organs^9–11^. However, few studies have investigated bone fracture healing at the single cell level and these studies have focused on cultured cells or late stages of bone repair^3,12^. Therefore, a complete dataset of periosteum and bone regeneration is lacking and is essential to decipher the mechanisms of pSSPC activation and regulation. Here, we created a single-nuclei atlas of the uninjured periosteum and its response to bone fracture. We generated single-nuclei RNAseq (snRNAseq) datasets from the uninjured periosteum and from the periosteum and hematoma/callus at days 3, 5 and 7 post-tibial fracture. Our analyses thoroughly describe the heterogeneity of the periosteum at steady state and the steps of pSSPC activation and differentiation after injury. We show that pSSPCs represent a single population localized in the fibrous layer of the periosteum. Periosteal SSPCs can provide osteoblasts and chondrocytes for bone repair by first generating a common injury-induced fibrogenic cell (IIFC) population that can then engage into osteogenesis and chondrogenesis. We identified the gene networks regulating pSSPC fate after injury and IIFCs as the main population producing paracrine factors mediating the initiation of bone healing.

## Results

### Heterogeneity of the periosteum at steady state

To investigate the heterogeneity of the periosteum at steady-state, we performed snRNAseq of the periosteum of wild-type mice (Fig. 1A-B). Single nuclei transcriptomics was previously shown to provide results equivalent to single cell transcriptomics, but with better cell type representation and reduced digestion-induced stress response^13–16^. After filtering, we obtained 1189 nuclei, corresponding to 8 cell populations: SSPC (expressing *Pi16*), fibroblasts (expressing *Pdgfra*), osteogenic cells (expressing *Runx2*), Schwann cells (expressing *Mpz*), pericytes/SMC (expressing *Tagln*), immune cells (expressing *Ptprc*), adipocytes (expressing *Lpl*), and endothelial cells (ECs, expressing *Pecam1*) (Fig. 1C-D, Fig. 1 – Supplementary Figure 1A). We performed in depth analyses of the SSPC, fibroblast and osteogenic cell populations. Subset analyses of clusters 0 to 5 identified 5 distinct SSPC/fibroblast populations expressing *Pdgfra* and *Prrx1*: *Pi16^+^ Ly6a* (SCA1)*^+^* cells (cluster 0), *Csmd1^+^* cells (cluster 1)*, Hsd11b1^+^* cells (cluster 2)*, Cldn1^+^* cells (cluster 3), and *Luzp2^+^* cells (cluster 4) (Fig. 2A-B). Cluster 0 is the only cell cluster containing cells expressing *Pi16*^+^ and stemness markers including *Ly6a* (SCA1)*, Dpp4,* and *Cd34* (Fig. 2C-D). Cytotrace scoring identified *Pi16*^+^ cells as the population in the most undifferentiated state (Fig. 2E). We performed in vitro CFU assays with sorted GFP^+^SCA1^+^ and GFP^+^SCA1^-^ cells isolated from the periosteum of *Prx1^Cre^; R26^mTmG^* mice, as *Prx1* labels all SSPCs contributing to the callus formation including *Pi16*^+^ cells (Fig 2D) ^1^. Prx1-GFP^+^ SCA1^+^ cells showed higher CFU potential compared to GFP^+^SCA1^-^ cells, confirming their stem/progenitor property (Fig 2F-G). Then, we grafted Prx1-GFP^+^ SCA1^+^ et Prx1-GFP^+^ SCA1^-^ periosteal cells at the fracture site of wild-type mice. Only GFP^+^SCA1^+^ cells formed cartilage after fracture indicating that GFP^+^SCA1^+^ cells encompass periosteal SSPCs with osteochondrogenic potential (Fig 2H). We explored the expression of other known markers of periosteal SSPCs, including *Ctsk*, *Acta2* (αSMA), *Gli1*, and *Mx1*, but no marker was fully specific to one cell cluster (Fig. 2 – Supplementary Fig. 1) ^1,2,5–7,17–21^.

**Figure 1:**
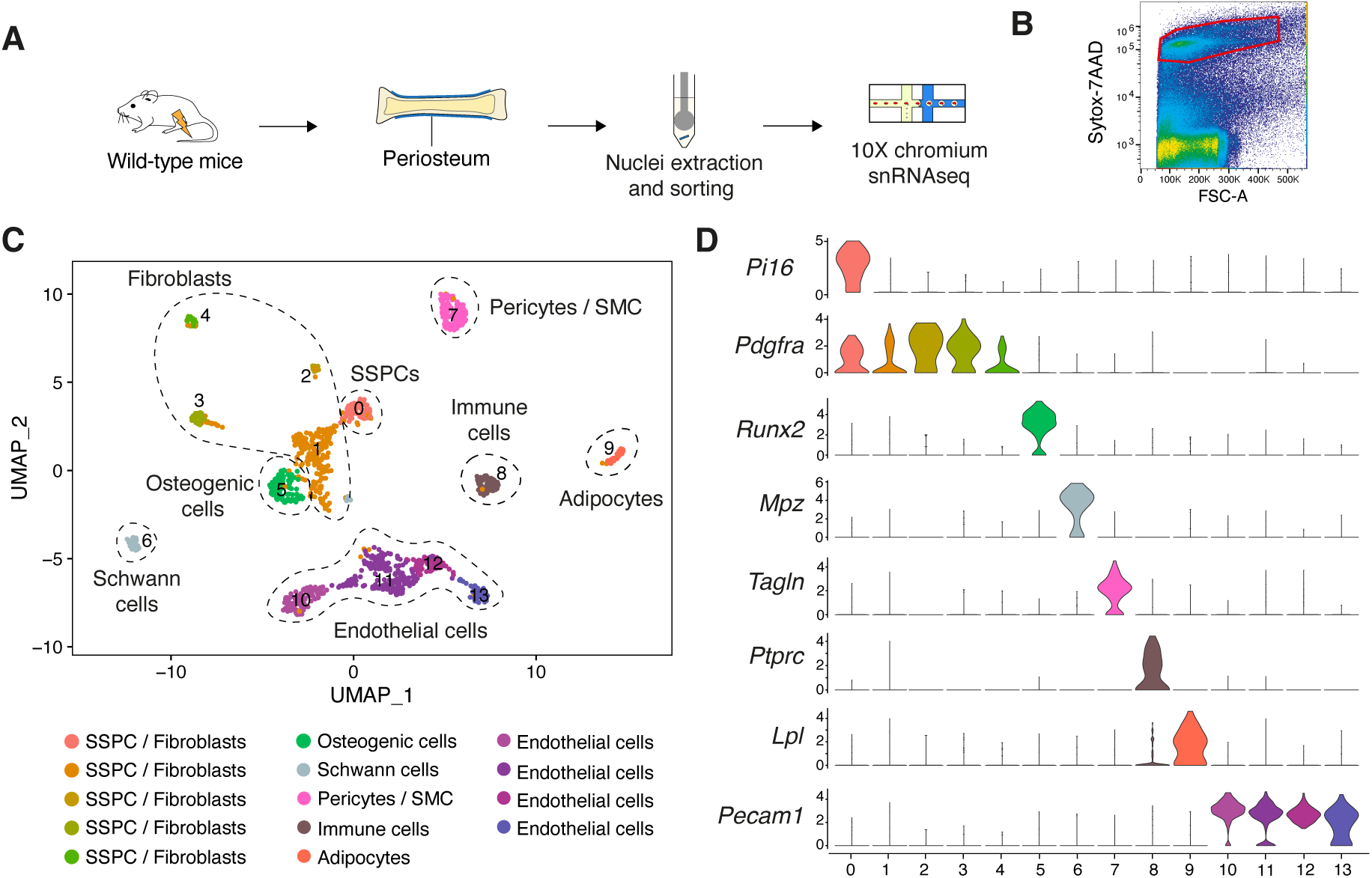
Heterogeneity of the periosteum at steady state. **A.** Experimental design. Nuclei were extracted from the periosteum of uninjured tibia and processed for single-nuclei RNAseq. **B.** Sorting strategy of nuclei stained with Sytox-7AAD for snRNAseq. Sorted nuclei are delimited by a red box. **C.** UMAP projection of color-coded clustering of the uninjured periosteum dataset. Six populations are identified and delimited by black dashed lines. **D.** Violin plots of key marker genes of the different cell populations.

**Figure 2:**
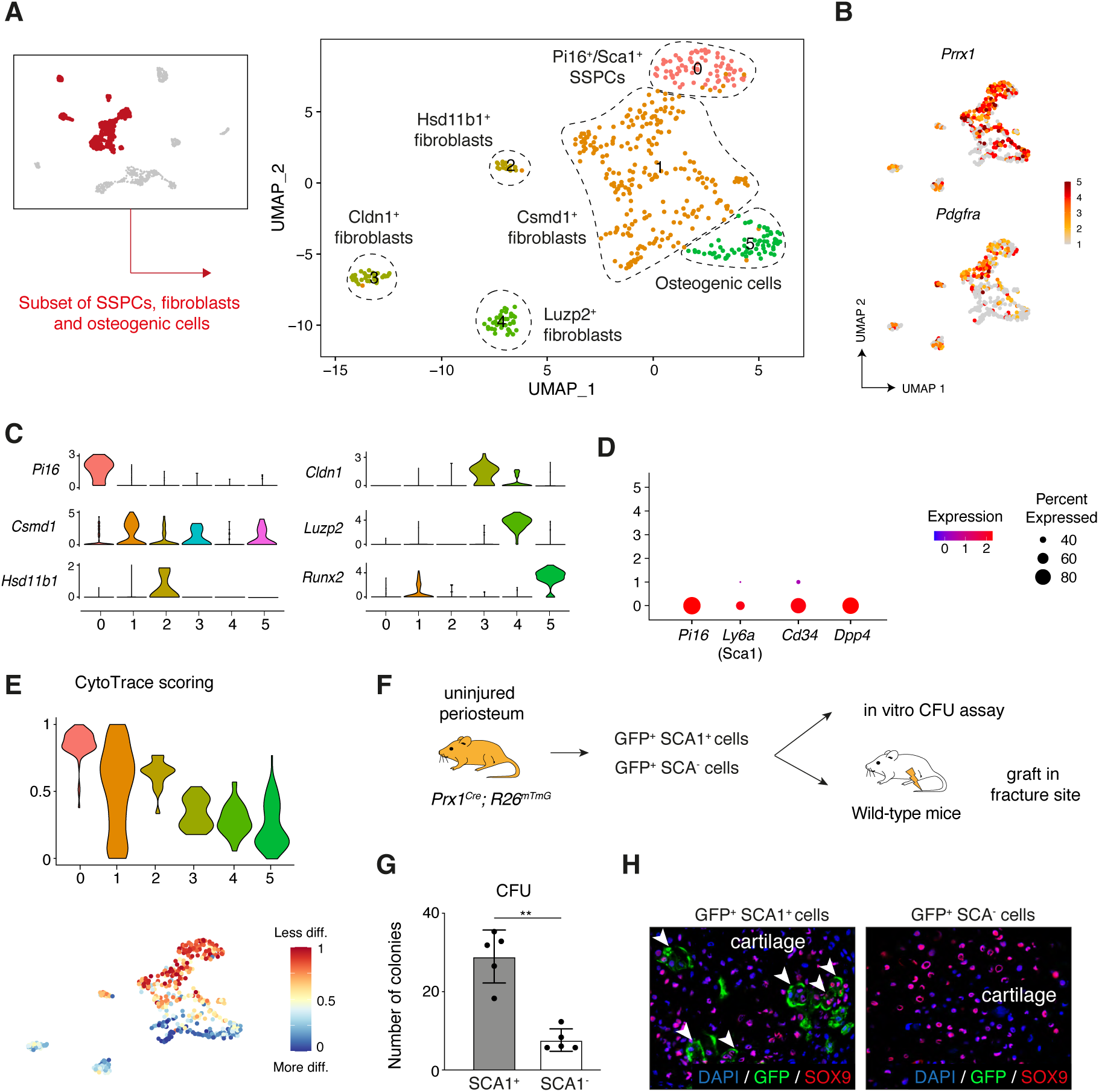
Identification of periosteal SSPCs in the intact periosteum. **A.** UMAP projection of color-coded clustering of the subset of SSPC/fibroblasts. **B.** Feature plots of *Prrx1* and *Pdgfra* in the subset of SSPC/fibroblasts. **C.** Feature plots of key marker genes of the different cell populations. **D.** Dot plot of the stemness markers *Pi16, Ly6a* (SCA1*), Cd34,* and *Dpp4*. **E.** Violin and feature plots of Cytotrace scoring in the subset of SSPC/fibroblasts, showing that SCA1^+^ SSPCs (cluster 0) are the less differentiated cells in the dataset. **F**. Experimental design: GFP^+^ SCA1^+^ and GFP^+^ SCA1^-^ were isolated from uninjured tibia of *Prx1^Cre^; R26^mTmG^* mice and used for in vitro CFU assays or grafted at the fracture site of wild-type mice. **G**. In vitro CFU assay of murine periosteal Prx1-GFP^+^ SCA1^+^ and Prx1-GFP^+^ SCA1^-^ cells. **H**. High magnification of SOX9 immunofluorescence of callus section 14 days post-fracture showing that GFP^+^SCA1^+^ cells contribute to the callus (white arrowheads) while GFP^+^ SCA1^-^ cells are not contributing (n=3 per group).

### The fracture repair atlas

To investigate the periosteal response to bone fracture, we collected injured periosteum with hematoma or callus at days 3, 5 and 7 post-fracture, extracted the nuclei and processed them for snRNAseq (Fig. 3A). We combined the datasets with the uninjured periosteum from Figure 1 and obtained a total of 6213 nuclei after filtering. The combined dataset was composed of 25 clusters corresponding to 11 cell populations: SSPCs (expressing *Pi16*), injury-induced fibrogenic cells (IIFCs, expressing ECM-related genes including *Postn*), osteoblasts (expressing *Ibsp*), chondrocytes (expressing *Col2a1*), osteoclasts (expressing *Ctsk*), immune cells (expressing *Ptprc*), Schwann cells (expressing *Mpz*), endothelial cells (expressing *Pecam1*), pericytes (expressing *Rgs5*), SMCs (expressing *Tagln*) and adipocytes (expressing *Lpl*) (Fig. 3B-C, Fig 3 – Supplementary Fig. 1-2). Next, we observed the dynamics of the cell populations in response to bone fracture (Fig. 3D, Fig 3 – Supplementary Fig. 1B). After injury, the percentage of SSPCs was strongly decreased and the percentage of IIFCs progressively increased (Fig. 3D-E). The percentage of chondrocytes and osteoblasts increased from day 3 post-fracture. Immune cells were drastically increased at day 3 after injury, before progressively decreasing at days 5 and 7 post-fracture (Fig. 3E).

**Figure 3:**
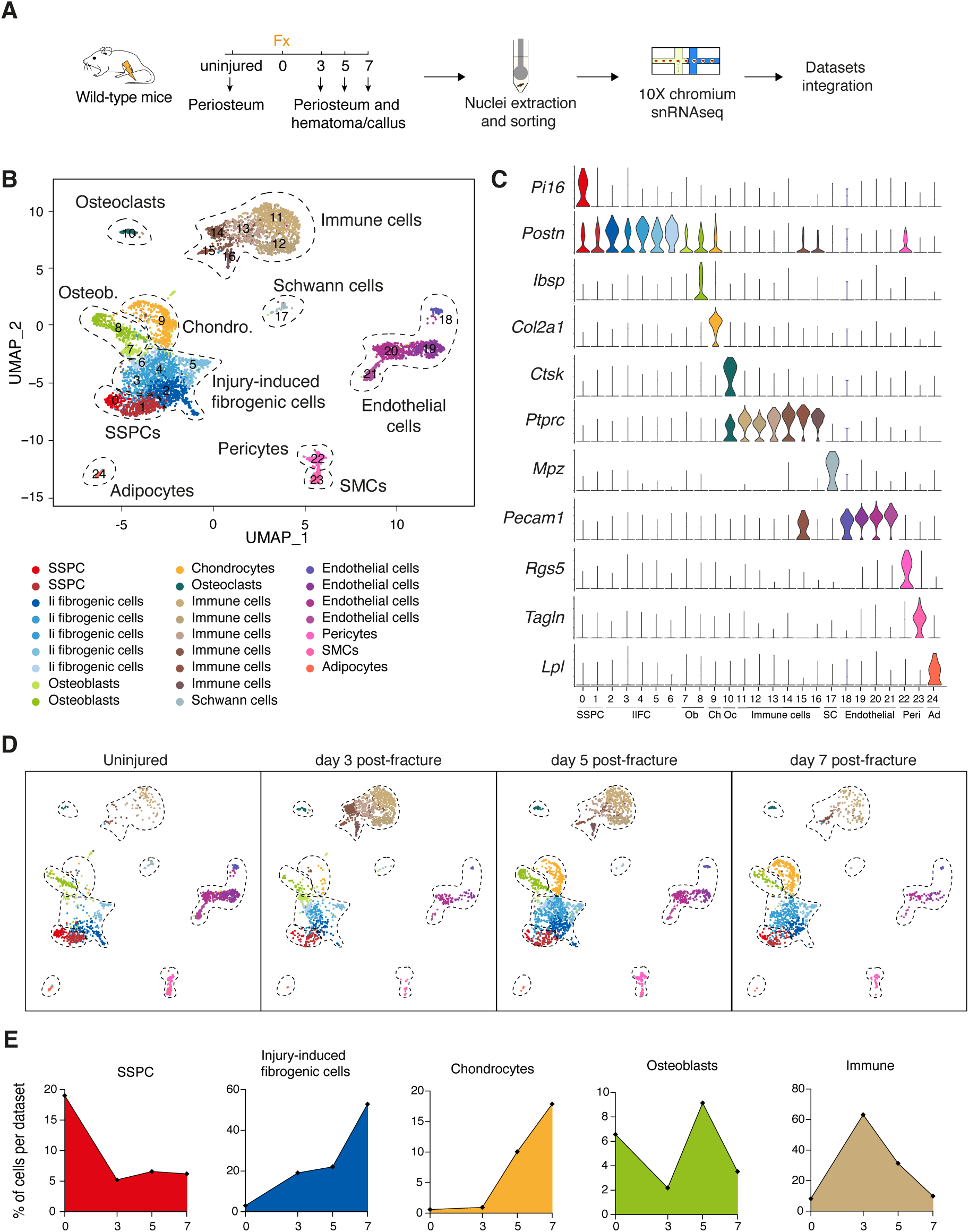
Periosteal response to fracture at single-nuclei resolution. **A.** Experimental design. Nuclei were extracted from the periosteum of uninjured tibia of wild-type mice and from the injured periosteum and hematoma/callus at days 3, 5 and 7 post-tibial fracture and processed for single-nuclei RNAseq. **B.** UMAP projection of color-coded clustering of the integration of uninjured, day 3, day 5 and day 7 datasets. Eleven populations are identified and delimited by black dashed lines. **C.** Violin plots of key marker genes of the different cell populations. **D.** UMAP projection of the combined dataset separated by time-point. **E.** Percentage of cells in SSPC, injury-induced fibrogenic cell, osteoblast, chondrocyte, and immune cell clusters in uninjured, day 3, day 5 and day 7 datasets.

### Spatial organization of the fracture callus

To evaluate the spatial distribution of the main cell populations identified in the snRNAseq datasets, we performed in situ immunofluorescence and RNAscope experiments on uninjured periosteum and days 3, 5 and 7 post-fracture hematoma/callus tissues. In the uninjured periosteum, we detected *Pi16*-expressing SSPCs, *Postn*-expressing cells, OSX^+^ osteoblasts, CD31^+^ endothelial cells and CD45^+^ immune cells (Fig 4A-B). *Pi16*-expressing SSPCs were located within the fibrous layer, while *Postn*-expressing cells were found in the cambium layer and corresponded to *Runx2*- expressing osteogenic cells (Fig. 4 – Supplementary Fig. 1A-C). Although *Postn* expression was weak in uninjured periosteum, *Postn* expression was strongly increased in response to fracture, specifically in IIFCs (Fig. 4 – Supplementary Fig. 1D-E). At day 3 post-fracture, we observed periosteal thickening and the formation of a fibrous hematoma (Fig. 4C). We did not detect *Pi16*- expressing SPPCs, consistent with the absence of cells expressing SSPC markers in the day 3 snRNAseq dataset compared to uninjured periosteum (Fig. 4 – Supplementary Figure 2). POSTN^+^ IIFCs and immune cells were the main populations present in hematoma and activated periosteum. Few IIFCs in the activated periosteum expressed SOX9 and OSX and only the periosteum was vascularized (Fig. 4D). At days 5 and 7 post-fracture, the callus was formed mainly of fibrotic tissue, new bone formed on the periosteal surface at the periphery of the callus and small cartilage islets were detected in the center of the callus near the periosteal surface (Fig. 4E-H). The fibrotic tissue contained mostly IIFCs, as well as immune and endothelial cells. We also observed SOX9^+^ and OSX^+^ cells in the fibrotic tissue, while the cartilage was solely composed of SOX9^+^ chondrocytes. OSX^+^ osteoblasts were the main cell population detected in the new bone at the periosteal surface. We also observed a progressive reduction of POSTN^+^ cells and immune cells from day 5 to 7 and increased vascularization in the newly formed bone (Fig. 4E-H).

**Figure 4:**
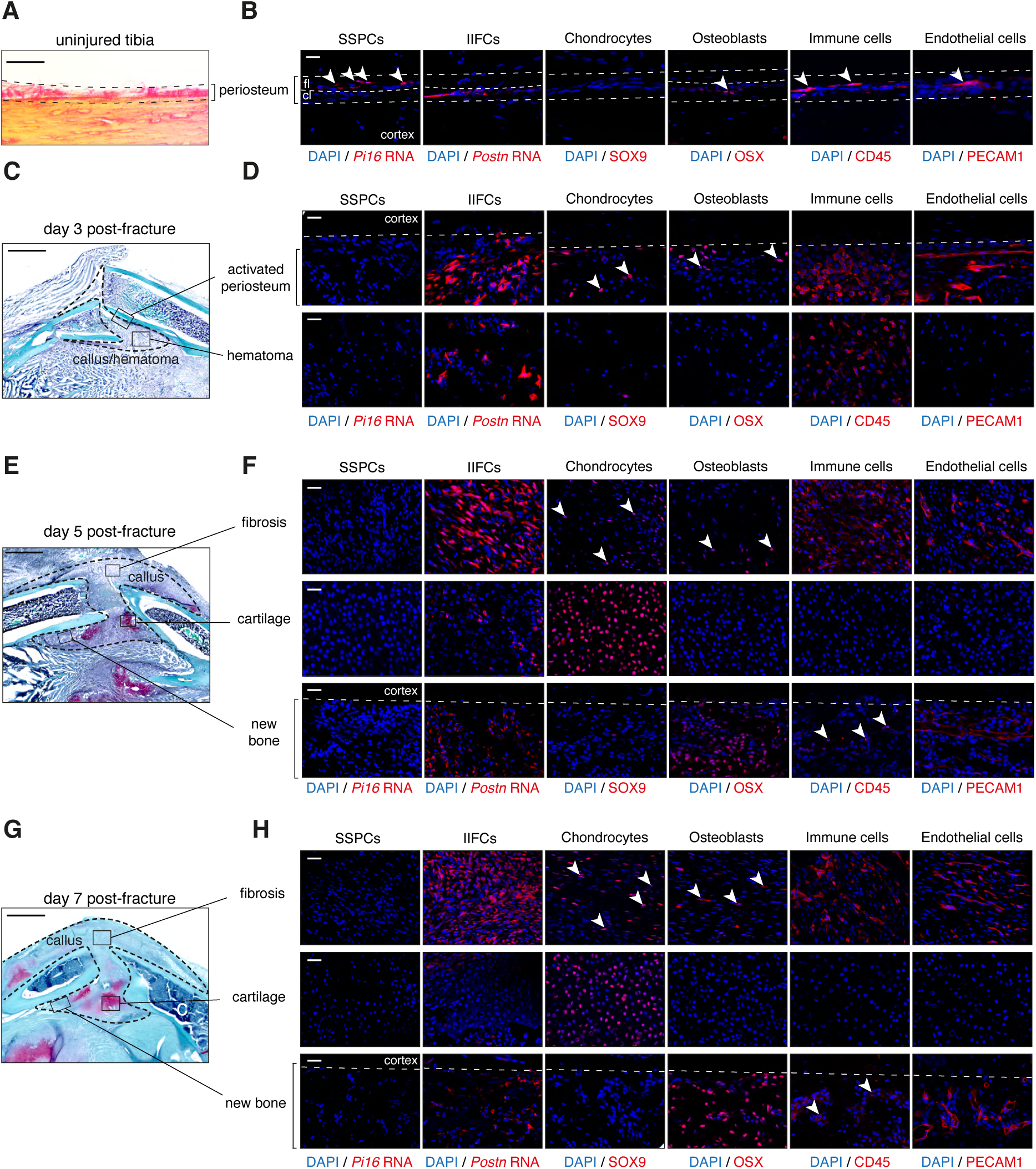
Cellular organization of the fracture callus. **A.** Picrosirius staining of the uninjured periosteum. **B**. Immunofluorescence and RNAscope on adjacent sections show the presence of SSPCs (*Pi16*-expressing cells) in the fibrous layer (fl), *Postn*-expressing cells in the cambium layer (cl), OSX^+^ osteoblasts, immune cells (CD45^+^) and endothelial cells (PECAM1^+^) in the periosteum (n=3 per group). **C.** Safranin’O staining of longitudinal callus sections at day 3-post tibial fracture. **D**. Immunofluorescence and RNAscope on adjacent sections show absence of SSPCs (*Pi16*^+^), and presence of IIFCs (*Postn*^+^) and immune cells (CD45^+^) in the activated periosteum and hematoma at day 3-post fracture. Chondrocytes (SOX9^+^,white arrowhead), osteoblasts (OSX^+^,white arrowhead), immune cells (CD45^+^) and endothelial cells (PECAM1^+^) are detected in the activated periosteum (n=3 per group). **E.** Safranin’O staining of longitudinal callus sections at day 5-post tibial fracture. **F**. Immunofluorescence and RNAscope on adjacent sections show IIFCs (*Postn*^+^), chondrocytes (SOX9^+^, white arrowhead), osteoblasts (OSX^+^,white arrowhead), immune cells (CD45^+^) and endothelial cells (PECAM1^+^) in the fibrosis, chondrocytes (SOX9^+^) in the cartilage and osteoblasts (OSX^+^), immune cells (CD45^+^,white arrowhead) and endothelial cells (PECAM1^+^) in the new bone. (n=3 per group). **G.** Safranin’O staining of longitudinal callus sections at day 7-post tibial fracture. **H**. Immunofluorescence and RNAscope on adjacent sections show IIFCs (*Postn* ^+^), chondrocytes (SOX9^+^,white arrowhead), osteoblasts (OSX^+^,white arrowhead), immune cells (CD45^+^) and endothelial cells (PECAM1^+^) in the fibrosis, chondrocytes (SOX9^+^) in the cartilage and osteoblasts (OSX^+^), immune cells (CD45^+^,white arrowhead) and endothelial cells (PECAM1^+^) in the new bone (n=3 per group). Scale bars: A-B-E: 1mm, B-D-F: 100µm.

### Periosteal SSPCs differentiate via an injury-induced fibrogenic stage

To understand the differentiation and fate of pSSPCs after fracture, we analyzed the subset of SSPC, IIFC, chondrocyte and osteoblast clusters from the combined fracture dataset (Fig. 5A, Fig. 5 – Supplementary Figure 1). We performed pseudotime analyses to determine the differentiation trajectories, defining the starting point in the pSSPC population, where the uninjured and undifferentiated cells cluster. We identified that pSSPCs differentiate in 3 stages starting from the pSSPC population (expressing *Ly6a*, *Pi16* and *Cd34*), predominant in uninjured dataset (Fig. 5B- C). Periosteal SSPCs then transition through an injury-induced fibrogenic stage predominant at days 3 and 5 post-injury. In this intermediate fibrogenic stage, IIFCs express high levels of extracellular matrix genes, such as *Postn*, *Aspn* and collagens. Subsequently, IIFCs differentiate into chondrocytes (expressing *Acan*, *Col2a1* and *Sox9*) or osteoblasts (expressing *Sp7*, *Alpl* and *Ibsp*), both predominant at days 5 and 7 (Fig. 5B-C, Table S1). We observed a parallel between pseudotime and the timepoints of the dataset, confirming that the differentiation trajectory follows the timing of cell differentiation (Fig. 5D). These results show that pSSPCs respond to fracture via an injury-induced fibrogenic stage common to chondrogenesis and osteogenesis and independent of their final fate (Fig. 5E). To visualize the transition of IIFCs towards chondrocytes and osteoblasts in the fracture callus, we performed co-immunofluorescence on day 5 post-fracture hematoma/callus. We observed a progressive increase in SOX9 and OSX signals in IIFCs at the fibrosis-to-cartilage and fibrosis-to-bone transition zones respectively (Fig 6A). To functionally validate the steps of pSSPC activation, we isolated SCA1^+^ GFP^+^ pSSPCs from *Prx1^Cre^; R26^mTmG^* mice, excluding endothelial cells (SCA1+GFP-) and pericytes (SCA1-GFP+), and grafted them at the fracture site of wild-type hosts (Fig. 6B, Figure 6 – Supplementary Figure 1). We observed that grafted GFP^+^ pSSPCs formed POSTN^+^ IIFCs at day 5 post-fracture (Fig 6B). Then, we isolated IIFCs, that correspond to GFP^+^ CD146^-^ cells from the day 3 post-fracture callus of *Prx1^Cre^; R26^mTmG^* mice without contamination by pericytes (GFP^+^CD146^+^ cells) (Fig. 6C, Figure 6 – Supplementary Figure 1). We grafted the GFP+ IIFCs at the fracture site of wild-type hosts and showed that grafted cells formed bone and cartilage at day 14 post-fracture. These results confirmed that pSSPCs first become IIFCs that differentiate into osteoblasts and chondrocytes.

**Figure 5:**
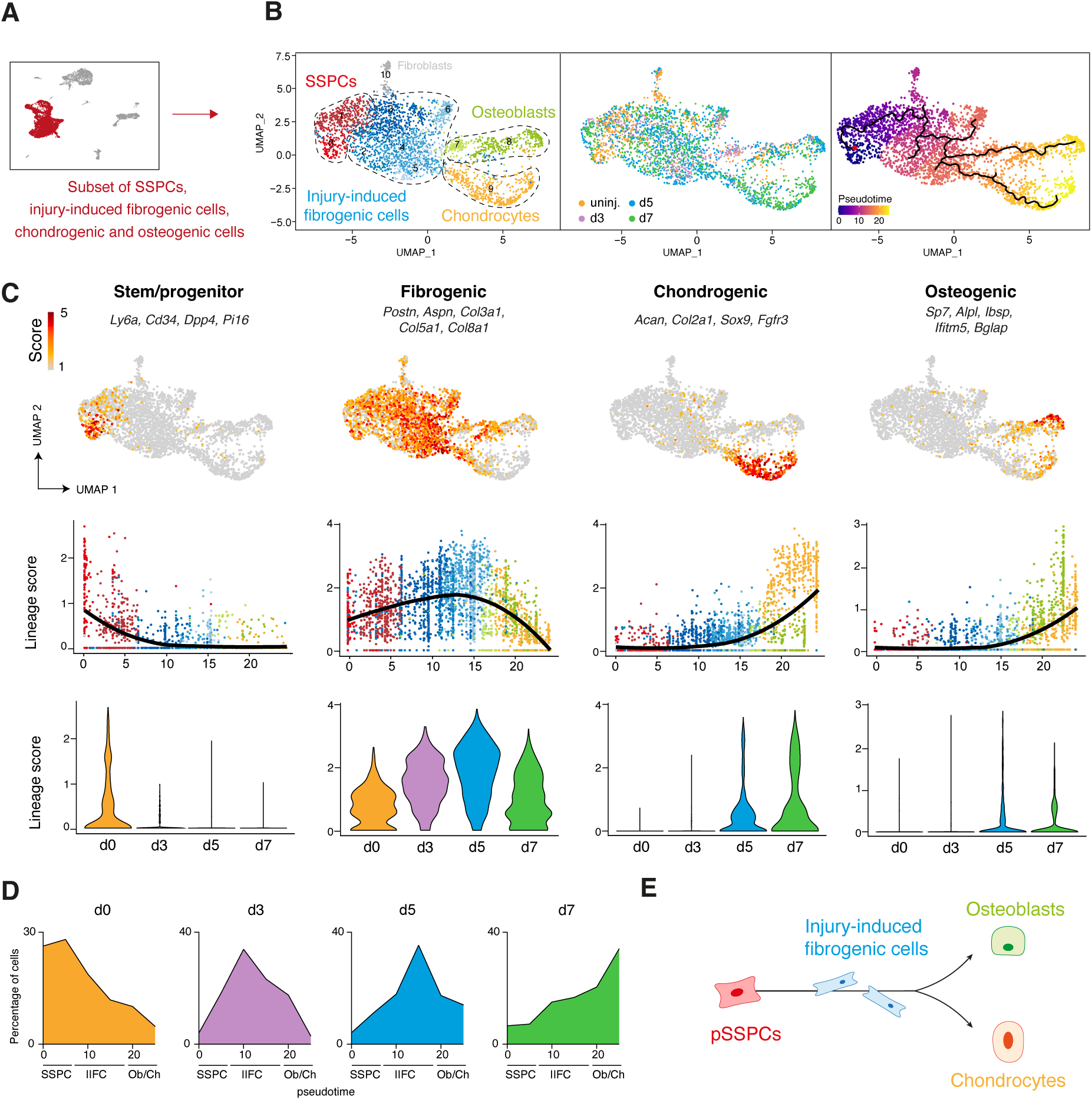
Periosteal SSPCs activate through a common fibrogenic state prior to undergoing osteogenesis or chondrogenesis. **A.** SSPCs, injury-induced fibrogenic cells (IIFCs), chondrocytes and osteoblasts from integrated uninjured, day 3, day 5 and day 7 post-fracture samples were extracted for a subset analysis. **B.** UMAP projection of color-coded clustering (left), color-coded sampling (middle) and monocle pseudotime trajectory (right) of the subset dataset. The four populations are delimited by black dashed lines. **C.** (top) Feature plots of the stem/progenitor, fibrogenic, chondrogenic and osteogenic lineage scores (middle) Scatter plot of the lineage scores along pseudotime. (bottom) Violin plot of the lineage score per time point. **D.** Distribution of the cells along the pseudotime per timepoint. **E.** Schematic representation of the activation trajectory of pSSPCs after fracture.

**Figure 6:**
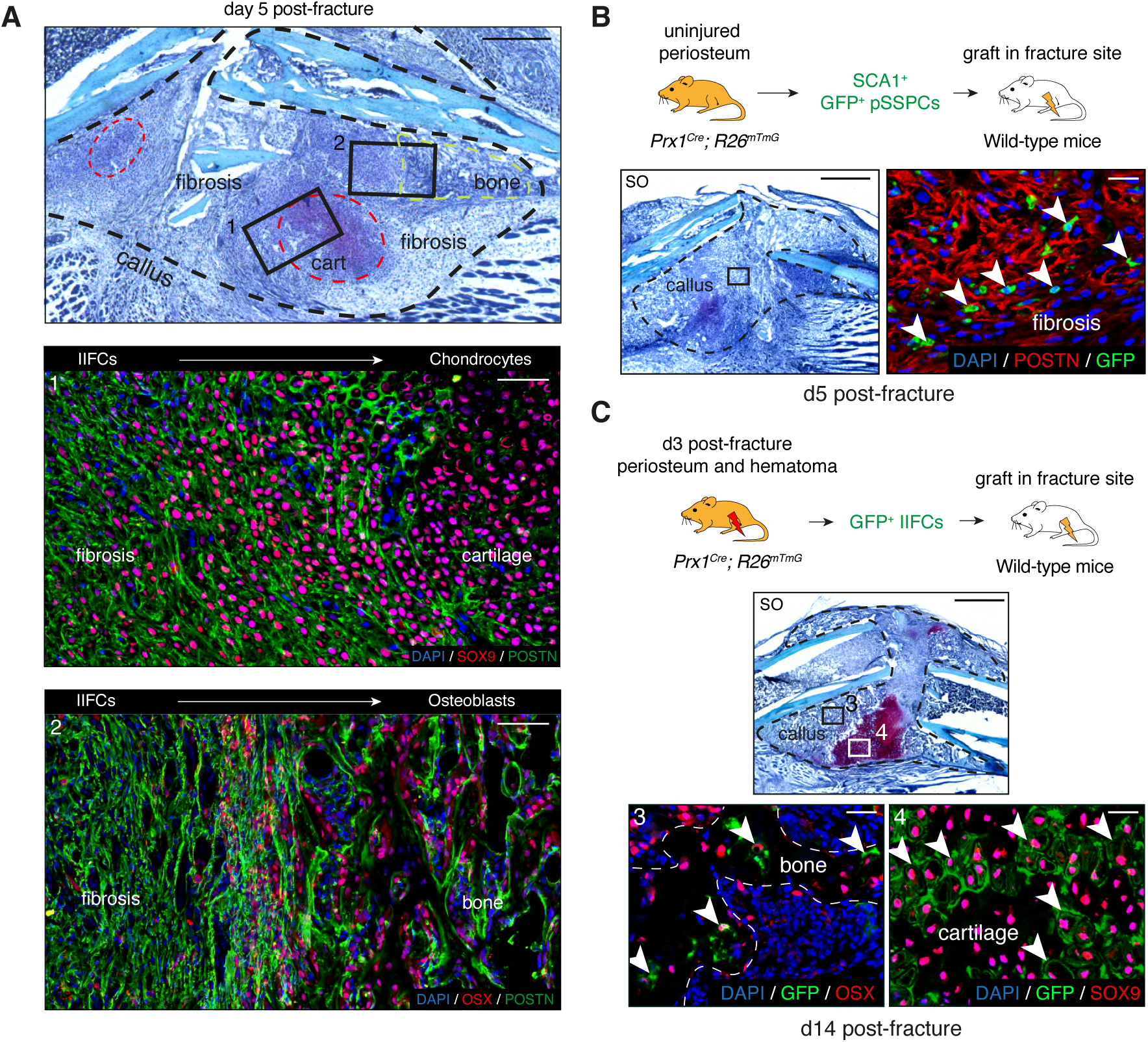
In vivo validation of pSSPC activation trajectory. **A**. (Top) Representative Safranin’O staining on longitudinal sections of the hematoma/callus at day 5 post-fracture. The callus is composed of fibrosis, cartilage (red dashed line) and bone (green dashed line). (Middle, box 1) Immunofluorescence on adjacent section shows decreased expression of POSTN (green) and increased expression of SOX9 (red) in the fibrosis-to-cartilage transition zone. (Bottom, box 2) Immunofluorescence on adjacent section shows decreased expression of POSTN (green) and increased expression of OSX (red) in the fibrosis-to-bone transition zone (n=3 per group). **B.** Experimental design: GFP^+^ SCA1^+^ SSPCs were isolated from uninjured tibia of *Prx1^Cre^; R26^mTmG^* mice and grafted at the fracture site of wild-type mice. Safranin’O staining of callus sections at day 5 post- fracture and high magnification of POSTN immunofluorescence of adjacent section showing that GFP^+^ cells contribute to the callus and that grafted SSPCs differentiate into POSTN^+^ IIFCS (white arrowheads) (n=4 per group). **C.** Experimental design: GFP^+^ IIFCs from periosteum and hematoma at day 3 post- fracture tibia were isolated from *Prx1^Cre^; R26^mTmG^* mice and grafted at the fracture site of wild-type mice. Safranin’O of callus sections at day 14 post-fracture and high magnification of OSX and SOX9 immunofluorescence of adjacent sections showing that GFP^+^ cells contribute to the callus and that grafted IIFCs differentiate into OSX^+^ osteoblasts (box 3, white arrowheads) and SOX9^+^ chondrocytes (box 4, white arrowheads) (n=4 per group). Scale bars: Low magnification: A: 500µm, B-C: 1mm. High magnification: 100µm.

### Characterization of injury-induced fibrogenic cells

We performed in depth analyses of the newly identified IIFC population. Gene ontology (GO) analyses of upregulated genes in IIFCs (clusters 2 to 6) showed enrichment in GOs related to tissue development, extracellular matrix (ECM), and ossification (Fig. 7A). We identified several ECM-related genes specifically upregulated in IIFCs, including collagens (*Col3a1, Col5a1, Col8a1, Col12a1*), *Postn*, and *Aspn* (Fig. 7B-C, Fig. 7 – Supplementary Figure 1). We also identified GO terms related to cell signaling, migration, differentiation, and proliferation, indicating that this step corresponds to an injury response. Only a small subset of IIFCs undergo apoptosis, further supporting that IIFCs are maintained in the fracture environment giving rise to osteoblasts and chondrocytes (Fig. 7 – Supplementary Figure 2). To further understand the mechanisms regulating SSPC activation and fate after injury, we performed gene regulatory network (GRN) analyses on the subset of SSPCs, IIFCs, osteoblasts and chondrocytes using SCENIC package (Single Cell rEgulatory Network Inference and Clustering)^22^. We identified 280 activated regulons (transcription factor/TF and their target genes) in the subset dataset. We performed GRN-based tSNE clustering and identified SSPC, IIFC, chondrocyte and osteoblast populations (Fig. 7D-E, Fig. 7-Supplementary Figure 3A). Fibroblasts from uninjured periosteum (*Hsd11b1^+^, Cldn1^+^*and *Luzp2^+^* cells and corresponding to cluster 10 of Fig. 5B) clustered separately from the other populations, suggesting the absence of their contribution to bone healing. Analysis of the number of activated regulons per cell indicated that SSPCs are the most stable cell population (higher number of activated regulons), while IIFCs are the less stable population, confirming their transient state (Fig. 7F). We then investigated cell population specific regulons. SSPCs showed activated regulons linked to stemness, including Hoxa10 (16g), Klf4 (346g), Pitx1 (10g), and Mta3 (228g) ^23–25^, and immune response, including Stat6 (30g), Fiz1 (72g) and Stat5b (18g)^26–28^ (Fig. 7G). Osteoblasts and chondrocytes display cell specific activated regulons including Sp7 (18g) and Sox9 (77g) respectively. We identified 21 regulons that we named fibro-core and that were upregulated specifically in the IIFC population (Fig. 7H-I, Table S2). Several fibro-core regulons, such as Meis1 (1556g), Pbx1 (11g), Six1 (20g), and Pbx3 (188g) are known to be involved in cell differentiation during tissue development and repair^29–32^. Reactome pathway analysis showed that the most significant terms linked to IIFC-specific TFs are related to Notch signaling (Fig. 7J, Table S3). We confirmed that Notch signaling is increased in the IIFC stage (Fig. 7J), suggesting its involvement in the fibrogenic phase of bone repair.

**Figure 7:**
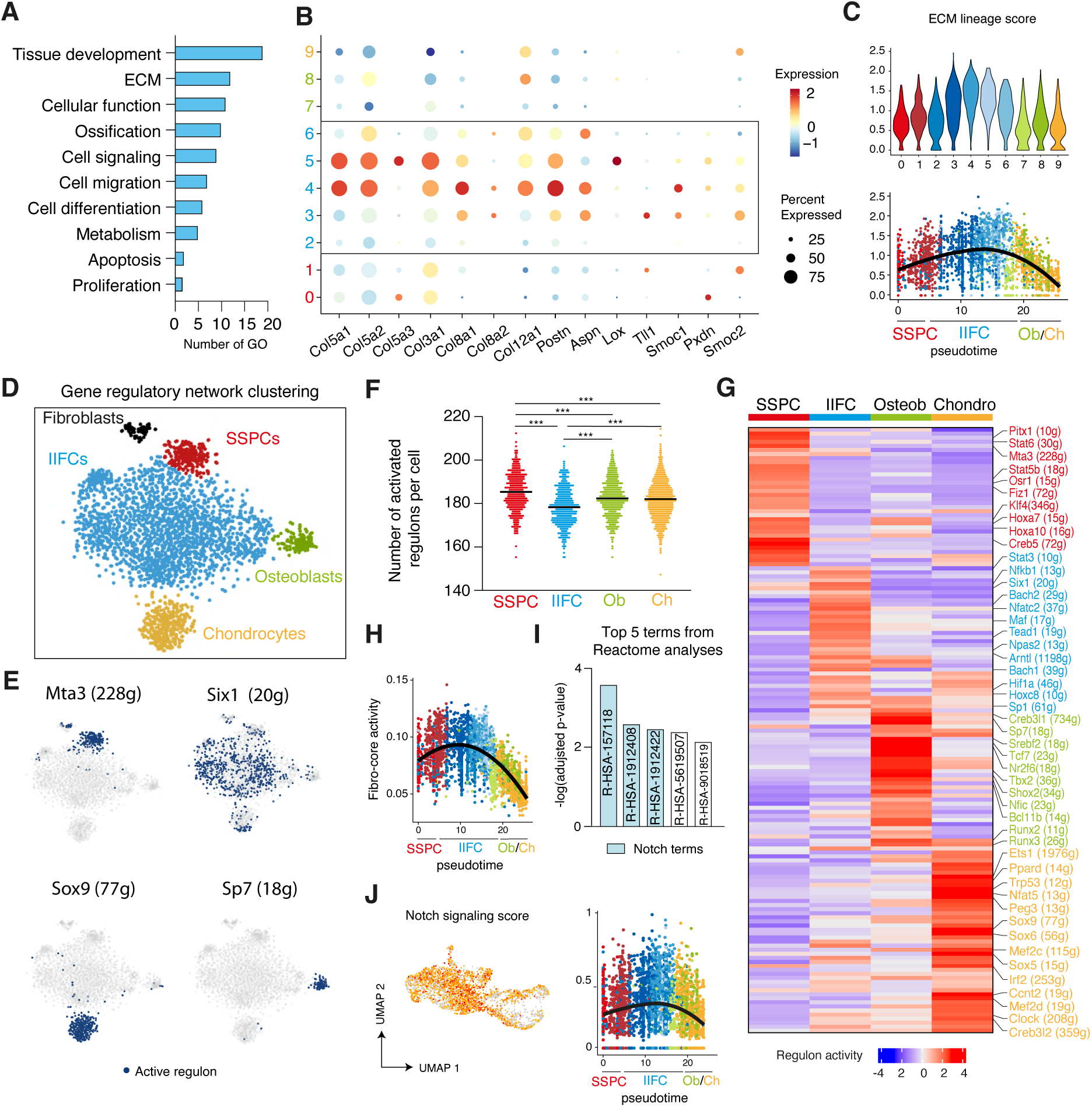
Characterization of injury-induced fibrogenic cells. **A.** Gene ontology analyses of upregulated genes in IIFCs (clusters 2 to 6 of UMAP clustering from Fig. 5). **B.** Dot plot of ECM genes in UMAP clustering from Fig. 5. **C.** Feature plot per cluster and scatter plot along pseudotime of the mean expression of ECM genes. **D.** Gene regulatory network (GRN)-based tSNE clustering of the subset of SSPCs, IIFCs, chondrocytes and osteoblasts. **E.** Activation of Mta3, Six1, Sox9 and Sp7 regulons in SSPCs, IIFCs, chondrocytes and osteoblasts. Blue dots mark cells with active regulon. **F.** Number of regulons activated per cell in the SSPC, IIFC, osteoblast (Ob) and chondrocyte (Ch) populations. Statistical differences were calculated using one-way ANOVA. ***: p-value < 0.001. **G.** Heatmap of activated regulons in SSPC, IIFC, osteoblast (osteob) and chondrocyte (chondro) populations. **H.** Scatter plot of the activity of the combined fibrogenic regulons along monocle pseudotime from Fig. 5. **I.** Reactome pathway analyses of the fibrogenic regulons shows that the 3 most significant terms are related to Notch signaling (blue). **J.** Feature plot in Seurat clustering and scatter plot along monocle pseudotime of the Notch signaling score.

### Distinct gene cores regulate the engagement of IIFCs in chondrogenesis and osteogenesis

We sought to identify the drivers of the transition of IIFCs to chondrocytes or osteoblasts. We identified 2 cores of regulons involved in chondrogenic differentiation. Chondro-core 1 is composed of 9 regulons specific to the transition of IIFCs to chondrocytes, including Maf (17g), Arntl (1198g), and Nfatc2 (37g) and chondro-core 2, composed of 14 regulons specific to differentiated chondrocytes (Fig. 8A, Fig. 8-Supplementary Figure 1A). Chondro-core 1 regulons are known to be regulators of the circadian clock (Npas2, Arntl) or of the T/B-cell receptor cellular response (Bach2, Nfact2, Nfkb1). STRING network analysis showed that TFs from the chondro-core 1 are interacting between each other and are at the center of the interactions between the chondro-core 2 TFs, including Sox9, Trp53 and Mef2c (Fig. 8B, Fig. 8-Supplementary Figure 1B). We observed that chondro-core 1 is only transiently activated when IIFCs are engaging into chondrogenesis and precedes the activation of chondro-core 2 (Fig. 8C-D). Chondro-core 1 activity was high in early differentiated chondrocytes (low *Acan* expression) and progressively reduced as chondrocytes undergo differentiation, while chondro-core 2 activity was gradually increased as chondrocytes differentiate (Fig. 8D). This suggests that transient activation of the chondro-core 1 allows the transition of IIFCs into chondrocytes. Then, we investigated the osteogenic commitment of IIFCs. We identified 8 regulons forming the osteo-core and activated in IIFCs transitioning to osteoblasts, such as Tcf7 (23g), Bcl11b (14g), and Tbx2 (36g) (Fig. 8E, Fig. 8-Supplementary Figure 1C). STRING network analysis showed that the genes with the strongest interaction with the osteo-core TFs are mostly related to Wnt signaling (Fig. 8F). This reveals the role of Wnt signaling in this transition from early fibrogenic activation of pSSPCs to osteogenic differentiation during bone repair. We calculated the osteo-core activity and observed that it is gradually increased and maintained in osteogenic cells, showing that the osteo-core is required for the transition and maturation of IIFC into osteoblasts (Fig 8G-H). Overall, we identified distinct cores of regulons with distinct dynamics driving the transition of IIFCs into chondrocytes and osteoblasts.

**Figure 8:**
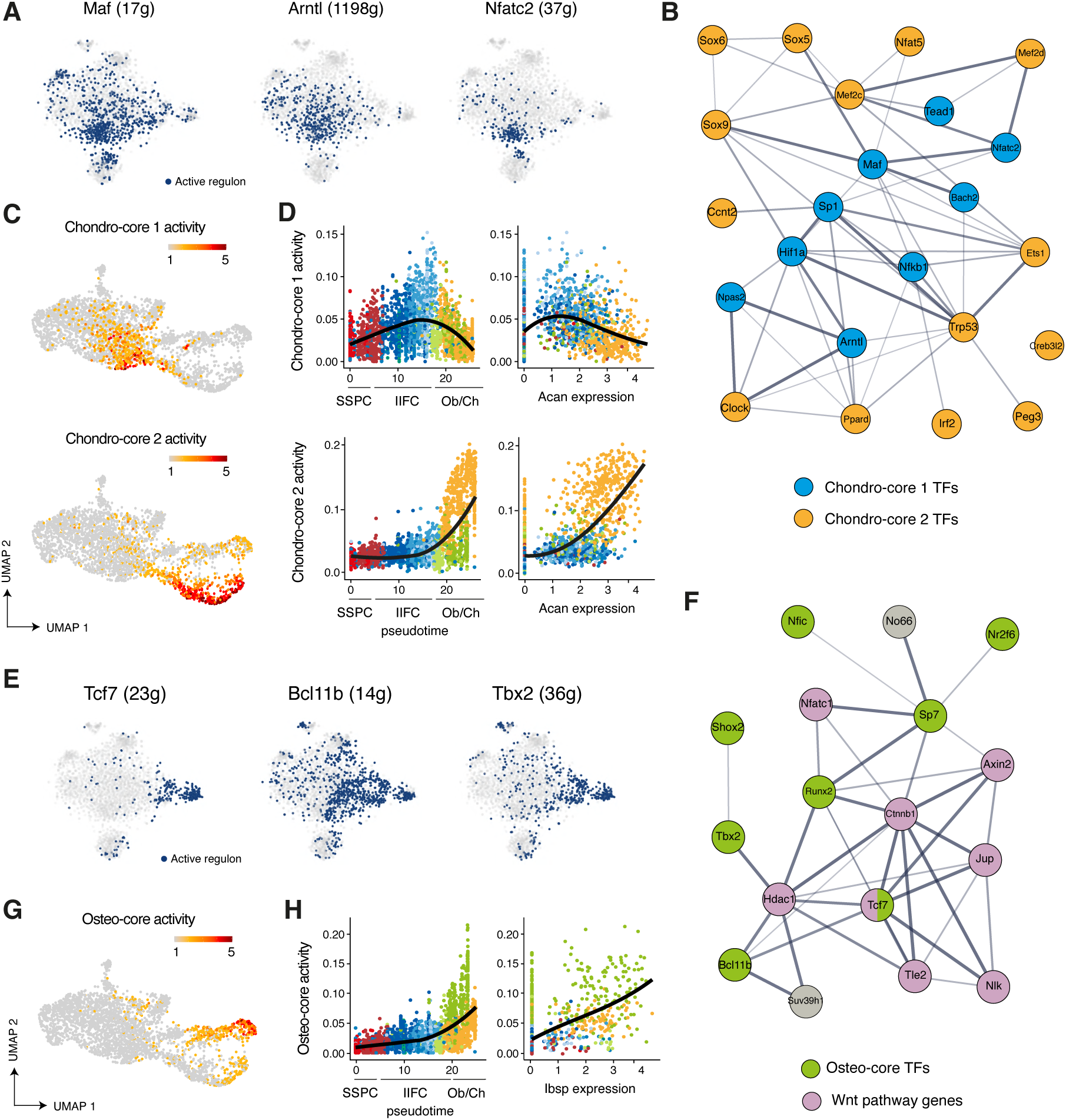
Gene regulatory network analyses identify gene cores driving fibrogenic to chondrogenic and osteogenic transitions. **A.** Activation of Maf, Arntl, and Nfatc2 regulons in SSPCs, IIFCs, chondrocytes and osteoblasts. **B.** STRING interaction network of the chondro-core 1 and 2 transcription factors (blue and orange respectively). **C.** Feature plot of chondro-core 1 (top) and chondro-core 2 (bottom) activities in SSPCs, IIFCs, chondrocytes and osteoblasts in Seurat UMAP from Fig. 5. **D.** Scatter plot of chondro-core 1 (top) and chondro-core 2 (bottom) activities along monocle pseudotime and *Acan* expressing. **E.** Activation of Tcf7, Bclb11b and Tbx2 regulons in SSPCs, IIFCs, chondrocytes and osteoblasts. **F**. STRING interaction network of the osteo-core transcription factors (green) and their related genes shows that most of osteo-core related genes are involved in Wnt pathway (purple). **G.** Feature plot of the osteo- core activity in SSPCs, IIFCs, chondrocytes and osteoblasts in Seurat UMAP from Fig. 5. **H.** Scatter plot of osteo-core activity along monocle pseudotime and *Ibsp* expressing.

### IIFCs mediate paracrine interactions during bone repair

Paracrine cell interactions are crucial drivers of tissue regeneration and stem cell activation. To identify key cell interactions during bone repair, we performed cell interaction analyses using CellChat package^33^. We observed that IIFCs are one of the predominant sources of outgoing signals during bone repair, and are also important receivers of signals, suggesting their central role in mediating cell interactions after fracture (Fig. 9A-B). Endothelial cells were mostly receiving signaling, while chondrocytes, osteoblasts and most immune cells exhibited reduced interactions with the other cell types in the fracture environment. IIFCs interact with all cell populations in the fracture environment, but the strongest interactions were with SSPCs and IIFCs (Fig. 9-Supplementary Figure 1A). CellChat analyses of the subset of SSPCs, IIFCs, osteoblasts and chondrocytes confirmed that IIFCs are a major source and receiver population of paracrine signals after fracture (Fig. 9-Supplementary Figure 1B). We then analyzed the main secreted factors from IIFCs. IIFCs secreted periostin (*Postn*), BMPs (*Bmp5*), pleiotropin/PTN (*Ptn*), TGFβs (*Tgfb2*, *Tgfb3*), PDGFs (*Pdgfc*, *Pdgfd*) and angiopoietin-likes/ANGPTLs (*Angplt2, Angplt4*) (Fig. 9C-D, Fig. 9- Supplementary Figure 1C). We observed differences in the dynamics of these factors, as some of them peak at day 3 post-fracture such as TGFβ, while others peak at day 5 such as BMP, POSTN, PTN and ANGPLT (Fig. 9E). We assessed the dynamics of ligand and receptor expression and observed that ligand expression was increased during the IIFC phase and specific of the fracture response (Fig. 9F). Receptor expression was high in both SSPCs and IIFCs, and receptors were expressed from steady state, suggesting that SSPCs can receive signals from IIFCs. Analysis of SSPC incoming signals showed that IIFCs produce paracrine factors that can regulate SSPCs (Fig. 9G), indicating that they contribute to SSPC recruitment after fracture.

**Figure 9:**
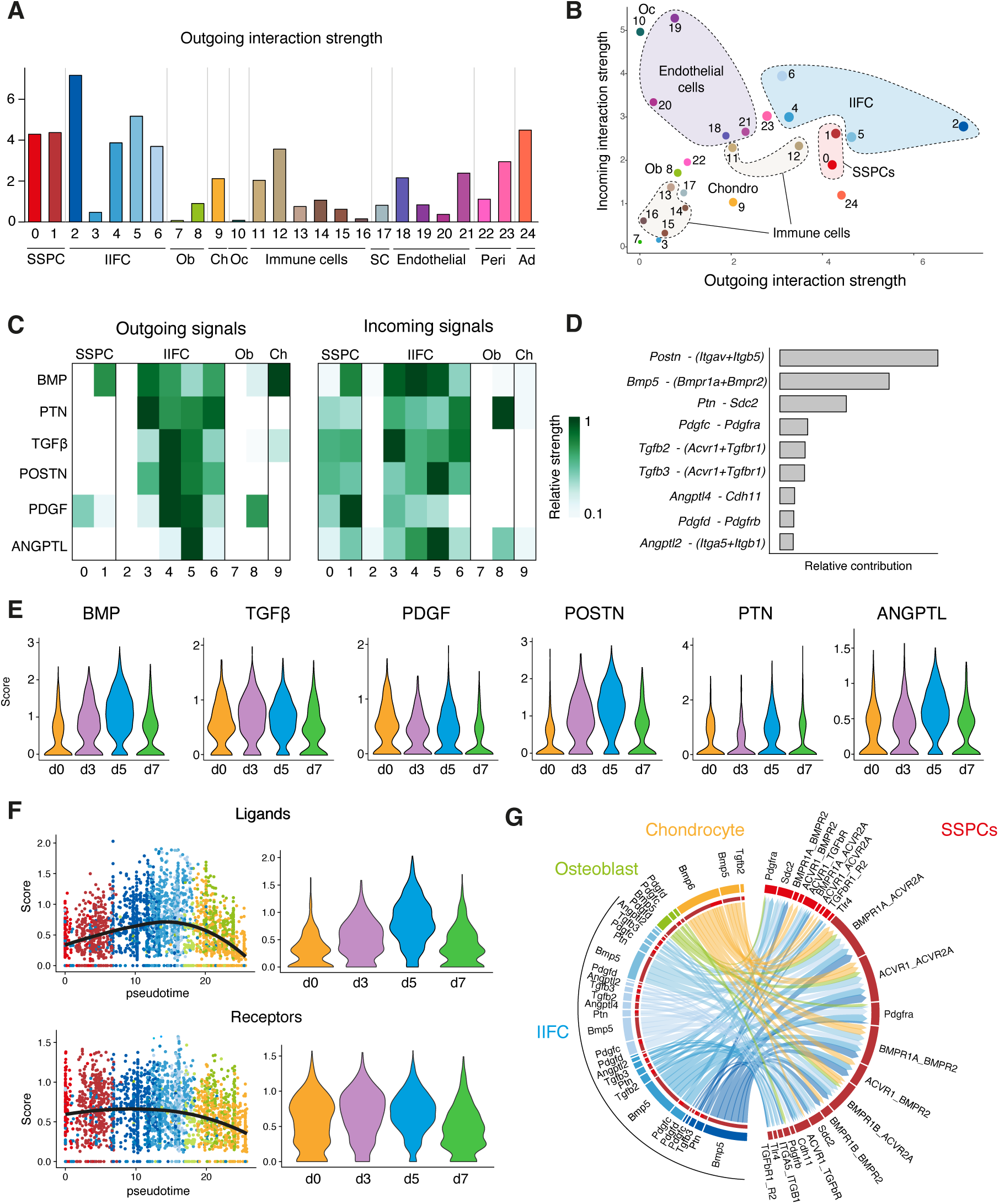
IIFCs are the main source of paracrine factors after fracture. **A.** Outgoing interaction strengths of the different cell populations of the fracture environment determined using CellChat package. **B.** Comparison of incoming and outgoing interaction strengths across SSPC, IIFC, chondrogenic and osteogenic populations. **C.** Outgoing and incoming signaling from and to SSPCs, IIFCs, chondrocytes and osteoblasts. **D.** Cell-cell interactions identified between SSPCs, IIFCs, chondrocytes and osteoblasts. **E.** Violin plots of the score of BMP, TGFβ, PDGF, POSTN, PTN and ANGPTL signaling per timepoint. **F.** Scatter plot along pseudotime and violin plot per time point of the mean expression of the ligand and receptors involved in signaling from IIFCs. **G.** Circle plot of the interactions between SSPCs, IIFCs, chondrocytes and osteoblasts, showing that most signals received by SSPCs are coming for IIFCs. Ob: osteoblasts, Oc: osteoclasts, Ch: chondrocytes, SC: Schwann cells, Ad: Adipocytes.

## Discussion

The periosteum is the main driver of bone regeneration. Yet the periosteum composition and its response to bone fracture are poorly described. Here, we used single-nuclei transcriptomics to understand the heterogeneity of the periosteum at steady state and the changes occurring within the periosteum after bone fracture. We developed a protocol to extract nuclei from freshly dissected periosteum, allowing us to capture their intact transcriptomic profile without enzymatic digestion- induced stress^13,34,35^. In addition, we performed snRNAseq without sorting specific populations, allowing the identification of all cell types located in the periosteum and the fracture environment from the early stages of repair. Our study provides the first complete fracture repair atlas, a key tool to understand bone regeneration.

First, we described the heterogeneity of the periosteum at steady state. While previous studies performed scRNAseq on sorted periosteal cell populations, our dataset uncovers the diversity of cell populations within the periosteum. We describe fibroblast populations, as well as tissue resident immune cells, adipocytes, and blood vessel/nerve-related cells. We identified one SSPC population of undifferentiated cells localized in the fibrous layer of the periosteum and expressing markers such as SCA1, *Dpp4* and *Pi16,* known to mark fibroblasts with stemness potential^36^. No markers previously described such as *Ctsk*, or *Gli1* were fully specific of one cell cluster and of the undifferentiated SSPC population, suggesting that these markers may label heterogeneous cell populations^2,5,6,19^.

After fracture, the composition of the periosteum changes drastically, with the appearance of IIFC and immune cells from day 3 post-fracture, and of chondrocytes and osteoblasts from day 5 post-fracture. Previous studies based on in vitro analyses and in vivo lineage tracing demonstrated that the periosteum displays the unique potential to form both bone and cartilage after fracture^1,3,5^. Yet, it was still unknown if the periosteum contains a bipotent SSPC population or several SSPC populations with distinct potentials after injury. Here, we show that pSSPCs respond to injury and form bone and cartilage via a single trajectory emerging from SCA1^+^ SSPCs. Periosteal SCA1^+^ SSPCs become IIFCs, a state marked by a decreased expression of stemness markers and a strong expression of extracellular matrix genes, such as *Postn* and *Aspn* and activation of Notch-related TFs. Thus, bone injury does not induce an expansion of SCA1^+^ SSPCs, rather a transition toward IIFCs that progressively expand and represent the main cell population within the fracture callus until day 7 post-fracture. Following this fibrogenic step, IIFCs do not undergo cell death but undergo either osteogenesis or chondrogenesis. This newly identified transient IIFC state represents the crossroad of bone regeneration and SSPC differentiation. In tissues such as skeletal muscle, an early transient fibrogenic response is also required for regeneration supporting muscle stem cell activation and differentiation^37^. During bone repair, this initial fibrogenic process is an integral part of the SSPC differentiation process, and a transitional step prior to osteogenesis and chondrogenesis.

GRN analyses identified TFs regulating SSPC response to fracture and involved in several cardinal signaling pathways including Notch and Wnt. Previous studies reported the role of Notch and Wnt in bone repair ^8,38–48^. Notch inactivation at early stages of repair leads to bone non-union while Notch inactivation in chondrocytes and osteoblasts does not significantly affect healing, correlating with our data showing that Notch is crucial during the IIFC phase, prior to osteochondral commitment^39^. Our results show that Wnt activation occurs after Notch activation and is specific to IIFCs engaging into osteogenesis, confirming its crucial role as a regulator of osteogenesis ^8,45–48^. In addition, we identified a chondro-core of 9 regulons transiently activated when IIFCs engage into chondrogenesis. Among these regulons, several are involved in the autonomous circadian clock, including *Npas2* and *Arntl* (Bmal1). Bmal1 was previously shown to be a regulator of cartilage differentiation in bone development and homeostasis^49^. Bmal1 inactivation leads to chondrocyte apoptosis and disruption of signaling involved in cartilage formation, including TGFβ, Ihh, HIF1α and NFAT^50–53^. This suggests a role of the circadian clock genes as key regulators of chondrogenesis during bone repair. IIFCs are also revealed as crucial paracrine regulators of the fracture environment. IIFCs secrete factors including BMPs, PDGFs, TGFβs, and POSTN, known to be required for successful bone healing^1,3,18,54^. These signals potentially regulate the interactions between IIFCs and all other cell types in the fracture environment, but primarily SSPCs and IIFCs themselves. Thus, IIFCs appear to contribute to the recruitment of SSPCs and promote their own maturation and differentiation.

Overall, our study provides a complete dataset of the early steps of bone regeneration. The newly identified SSPC activation pattern involves a coordinated temporal dynamic of cell phenotypes and signaling pathways after injury. IIFCs emerge as a transient cell population that plays essential roles in the initial steps of bone repair. Deeper understanding of IIFC regulation will be crucial as they represent the ideal target to enhance bone healing and potentially treat bone repair defects.

## Materials and Methods

### Mice

C57BL/6ScNj were obtained from Janvier Labs (France). *Prx1^Cre^*^55^ and *Rosa26-mtdTomato- mEGFP* (*R26^mTmG^*)^56^ were obtained from Jackson Laboratory (Bar Harbor, ME). ). All SSPCs, including pSSPCs, are marked by GFP in *Prx1^Cre;^* R26^mTmG^ mice. Mice were bred in animal facilities at IMRB, Creteil and kept in separated ventilated cages, in pathogen-controlled environment and ad libitum access to water and food. All procedures performed were approved by the Paris Est Creteil University Ethical Committee. Twelve-week-old males and females were mixed in experimental groups.

### Tibial fracture

Open non-stabilized tibial fractures were induced as previously described ^57^. Mice were anesthetized with an intraperitoneal injection of Ketamine (50 mg/mL) and Medetomidine (1 mg/kg) and received a subcutaneous injection of Buprenorphine (0.1 mg/kg) for analgesia. Mice were kept on a 37°C heating pad during anesthesia. The right hindlimb was shaved and sanitized. The skin was incised to expose the tibia and osteotomy was performed in the mid-diaphysis by cutting the bone. The wound was sutured, the mice were revived with an intraperitoneal injection of atipamezole (1 mg/kg) and received two additional analgesic injections in the 24 hours following surgery. Mice were sacrificed at 3, 5, 7 or 14 days post-fracture.

### Nuclei extraction

Nuclei extraction protocol was adapted from ^58,59^. We generated 4 datasets for this study: uninjured periosteum, periosteum and hematoma at days 3, 5 and 7 post-tibial fracture. The uninjured and day 3 post-fracture datasets were generated in duplicates. For uninjured periosteum, tibias from 4 mice were dissected free of muscle and surrounding tissues. The epiphyses were cut and the bone marrow flushed. The periosteum was scraped from the cortex using dissecting Chisel (10095-12, Fine Science Tools). For days 3, 5 and 7 post fracture, injured tibias from 4 to 9 mice were collected and the surrounded tissues were removed. The activated periosteum was scraped and collected with the hematoma. Collected tissues were minced and placed 5 min in ice-cold Nuclei Buffer (NUC101, Merck) before mechanical nuclei extraction using a glass douncer. Extraction was performed by 20 strokes of pestle A followed by 5-10 of pestle B. Nuclei suspension was filtered, centrifuged and resuspended in RNAse-free PBS (AM9624, ThermoFischer Scientific) with 2% Bovine Serum Albumin (A2153, Merck) and 0.2 U/µL RNAse inhibitor (3335399001, Roche). A second step of centrifugation was performed to reduce contamination by cytoplasmic RNA. Sytox™ AADvanced™ (S10349, ThermoFischer Scientific) was added (1/200) to label nuclei and Sytox-AAD+ nuclei were sorted using Sony SH800.

### Single nuclei RNA sequencing

The snRNA-seq libraries were generated using Chromium Single Cell Next GEM 3′ Library & Gel Bead Kit v.3.1 (10x Genomics) according to the manufacturer’s protocol. Briefly, 10 000 to 20 000 nuclei were loaded in the 10x Chromium Controller to generate single-nuclei gel-beads in emulsion. After reverse transcription, gel-beads in emulsion were disrupted. Barcoded complementary DNA was isolated and amplified by PCR. Following fragmentation, end repair and A-tailing, sample indexes were added during index PCR. The purified libraries were sequenced on a Novaseq (Illumina) with 28 cycles of read 1, 8 cycles of i7 index and 91 cycles of read 2. Sequencing data were processed using the Cell Ranger Count pipeline and reads were mapped on the mm10 reference mouse genome with intronic and exonic sequences.

### Filtering and clustering using Seurat

Single-nuclei RNAseq analyses were performed using Seurat v4.1.0 ^60,61^ and Rstudio v1.4.1717. Aligned snRNAseq datasets were filtered to retain only nuclei expressing between 200 et 5000 genes and expressing less than 2% of mitochondrial genes and 1.5% of ribosomal genes. Contamination from myogenic cells were removed from the analyses. After filtering, we obtained 1378 nuclei from uninjured periosteum, 1634 from day 3 post-fracture, 2089 from d5 post-fracture and 1112 from day 7 post-fracture. The replicates of the uninjured dataset were integrated using Seurat. The integrated dataset was regressed on cell cycle, mitochondrial and ribosomal content and clustering was performed using the first 15 principal components and a resolution of 0.5. SSPC/fibroblast and osteogenic cells were isolated and reclustered using the first 10 principal components and a resolution of 0.2. Uninjured, d3, d5 and d7 datasets were integrated using Seurat. The integrated dataset was regressed on cell cycle, mitochondrial and ribosomal content. Clustering was performed using the first 20 principal components and a resolution of 1.3. SSPC, IIFC, chondrogenic and osteogenic clusters from the integration were isolated to perform subset analysis. The subset was reclustered using the first 15 principal components and a resolution of 0.6.

### Pseudotime analysis using monocle3

Monocle3 v1.0.0 was used for pseudotime analysis ^62^. Single-cell trajectories were determined using monocle3 default parameters. The starting point of the pseudotime trajectory was determined as the cells from the uninjured dataset with the highest expression of stem/progenitor marker genes *(Ly6a, Cd34, Dpp4, Pi16)*. Pseudotime values were added in the Seurat object as metadata and used with the Seurat package.

### Differentiation state analysis using Cytrotrace

To assess the level of differentiation of the cell clusters, we performed analyses using CytoTrace with the default parameters. Cytotrace scoring was plotted in violin plot and on the Seurat UMAP clustering.

### Gene Ontology and Reactome and analyses

Reactome and GO analyses were performed using EnrichR ^63^. All significant GO terms from upregulated genes in clusters 2 to 6 of the subset of SSPCs, IIFCs, osteoblasts and chondrocytes were manually categorized. The 5 more significant terms of the Reactome analysis from the fibrogenic TFs of Fig. 6 are presented in Table S3.

### Single cell regulatory network inference using SCENIC

Single cell regulatory network inference and clustering (SCENIC) ^22^ was used to infer transcription factor (TF) networks active in SSPCs, IIFCs, osteoblasts and chondrocytes. Analysis was performed using recommended parameters using the packages SCENIC v1.3.1, AUCell v1.16.0, and RcisTarget v1.14 and the motif databases RcisTarget and GRNboost. SCENIC package was used to perform regulon based tSNE clustering and identified population specific regulons.

### Cell-cell interaction using CellChat

Cell communication analysis was performed using the R package CellChat^33^ with default parameters on the complete fracture combined dataset and on the subset of SSPCs, IIFCs, osteoblasts and chondrocytes.

### STRING network analyses

To assess protein-protein interaction network, we used the STRING v11.5 database^64^. To assess interaction in the chondro-core, we performed the analysis on the chondro-core and chondro-specific TFs identified in our analysis. For osteo-core analyses, we performed the analysis with osteo-core genes and the most significant interactions.

### Histology and immunofluorescence

Mouse samples were processed as previously described^57^. Tibias were collected and fixed in 4% PFA (sc-281692, CliniSciences) for 4 hours at 4°C. Then, samples were decalcified in 19% EDTA for 10 days (EU00084, Euromedex), cryoprotected in 30% sucrose (200-301-B, Euromedex) for 24h and embedded in OCT. Samples were sectioned in 10µm thick sections. Cryosections were defrosted and rehydrated in PBS. For Safranin-O staining, sections were stained with Weigert’s solution for 5 min, rinsed in running tap water for 3 min and stained with 0.02% Fast Green for 30 seconds (F7252, Merck), followed by 1% acetic acid for 30 seconds and Safranin’O solution for 45 min (S2255, Merck). For immunofluorescence, sections were incubated 1 hour at room temperature in 5% serum, 0.25% Triton PBS before incubation overnight at 4°C with the following antibodies: Rat monoclonal to mouse SCA1 (740450, BD Biosciences), Rabbit monoclonal to mouse SOX9 (ab185230, Abcam), Rabbit polyclonal to mouse Osterix/Sp7 (ab22552, Abcam), Goat polyclonal to mouse Periostin (AF2955, R&D Systems), Goat polyclonal to mouse PECAM1 (AF3628, Biotechne), Rat monoclonal to mouse CD45 (552848, BD Bioscience). Secondary antibody incubation was performed at room temperature for 1 hour. Slides were mounted with Fluoromount-G mounting medium with DAPI (00-4959-52, Life Technologies).

### RNAscope in situ hybridization

The expression of *Pi16* and *Postn* was visualized using the RNAscope® Multiplex Fluorescent Assay V2 (Biotechne). Tissues were processed as described above. Ten µm thick sections were cut and processed according to the manufacturer’s protocol: 15 min of post-fixation in 4% PFA, ethanol dehydration, 10 min of H2O2 treatment and incubation in ACD custom reagent for 30 minutes at 40°C. After hybridization and revelation, the sections were mounted under a glass coverslip with Prolong Gold Antifade (P10144, ThermoFischer).

### Tissue dissociation and cell sorting

#### Periosteal SSPC isolation

To isolate periosteal cells, uninjured tibias from *Prx1^Cre^; R26^mTmG^* mice were collected, and all surrounding soft tissues were carefully removed. Epiphyses were embedded in low melting agarose and tibias were placed for 30 at 37°C in digestion medium composed of PBS with 3mg/ml of Collagenase B (C6885, Merck), 4mg/ml of Dispase (D4693, Merck) and 100U/mL of DNAse I (WOLS02007, Serlabo, France). After digestion, tibias were removed and the suspension was filtered, centrifuged and resuspended.

#### Injury-induced fibrogenic cell isolation

The fracture hematoma and the activated periosteum were collected from *Prx1^Cre^; R26^mTmG^* mice 3 days post-fracture. Tissues were minced and digested at 37°C for 2 hours in DMEM (21063029, Life Technologies) with 1% Trypsin (15090046, Life Technologies) and 1% collagenase D (11088866001, Roche). Cells in suspension were removed every 20 min and the digestion medium was replaced. After 2 hours, the cell suspension was filtered, centrifuged and resuspended.

#### Cell sorting and transplantation

Digested cells were incubated 30 minutes with anti-SCA1-BV650 (BD Biosciences, 740450) or anti- CD146-BV605 (BD Biosciences, 740636). Cells were sorted using Influx Cell Sorter. Prior sorting, Sytox Blue (S34857, Thermo Fisher Scientific) was added to stain dead cells. We sorted living single GFP^+^ SCA1^+^ (pSSPCs excluding GFP^-^ endothelial cells and SCA1^-^ pericytes) and living single GFP^+^ SCA1^-^ cells from the uninjured periosteum of *Prx1^Cre^;Rosa^mTmG^* mice. For IIFCs, living single GFP^+^ CD146^-^ cells were sorted, to eliminate CD146^+^ pericytes. Cell transplantation was performed as described in ^57^. 30000 to 45000 sorted GFP^+^ cells were embedded in Tisseel Prima fibrin gel, composed of fibrinogen and thrombin (3400894252443, Baxter S.A.S, USA), according to manufacturer’s instructions. Briefly, the cells were resuspended in 15 μL of fibrin (diluted at 1/4), before adding 15 μL of thrombin (diluted at 1/4) and mixing. The pellet was then placed on ice for at least 15 min for polymerization. The cell pellet was transplanted at the fracture site of wild-type mice at the time of fracture.

### In vitro colony-forming Unit (CFU) assay

Prx1-GFP^+^ SCA1^+^ and Prx1-GFP^+^ SCA1^-^ periosteal cells were isolated from *Prx1^Cre^;Rosa^mTmG^*mice as described above. Sorted cells were plated at a cell density of 2000 cells per 6 well-plates in growth media consisting of MEMα supplemented with 20% FBS, 1% penicillin-streptomycin (Life Technology, Carlsbad, California), and 0.01% bFGF (R&D, Minneapolis, MN). The medium was changed every 2-3 days. After 11 days, plates were washed twice with PBS, fixed with methanol, stained with 0,5% crystal violet diluted in 20% methanol and colonies were manually counted.

### Statistical analyses

Statistical difference between the number of activated regulons per cell was determined using one way-ANOVA followed by a post-hoc test. A p-value < 0.001 is reported as ***.

### Resource availability

Single nuclei RNAseq datasets are deposited in the Gene Expression Omnibus (GSE234451). The integrated dataset was deposited on USCS Cell Browser for easy use: (https://fracture-repair-atlas.cells.ucsc.edu). This paper does not report original code. Further information and requests for resources and reagents should be directed to and will be fulfilled by the lead contact, Céline Colnot.

## Acknowledgements

This work was supported by ANR-18-CE14-0033, ANR-21-CE18-007-01, NIAMS R01 AR072707 and R01 AR081671 to C.C. S. Perrin, M. Ethel and C. Goachet were supported by a PhD fellowship from Paris Cité University, Univ Paris-Est Creteil and Fondation pour la Recherche Médicale respectively. We thank O. Ruckebusch and A. Guigan from the Flow Cytometry platform at IMRB Institute, O. Pellé from the Flow Cytometry platform at Imagine Institute and all the staff from the Imagine genomic core facility at Imagine Institute. We thank A. Julien, S. Protic and Y. Hachemi for technical assistance or advice.

## Author contributions

Conceptualization: S.P. and C.C. Methodology: S.P. and C.C. Formal analysis: S.P. and C-A.W. Investigation: S.P., M.E., V.B., C.G., M.L., F.C. and C.M. Resources: F.C. and M.M. Writing – Original Draft: S.P. and C.C. Visualization: S.P. Supervision: C.C. Project administration: C.C. Funding Acquisition: C.C

## Supplemental information

**Figure 1 – Supplementary Figure 1A:**
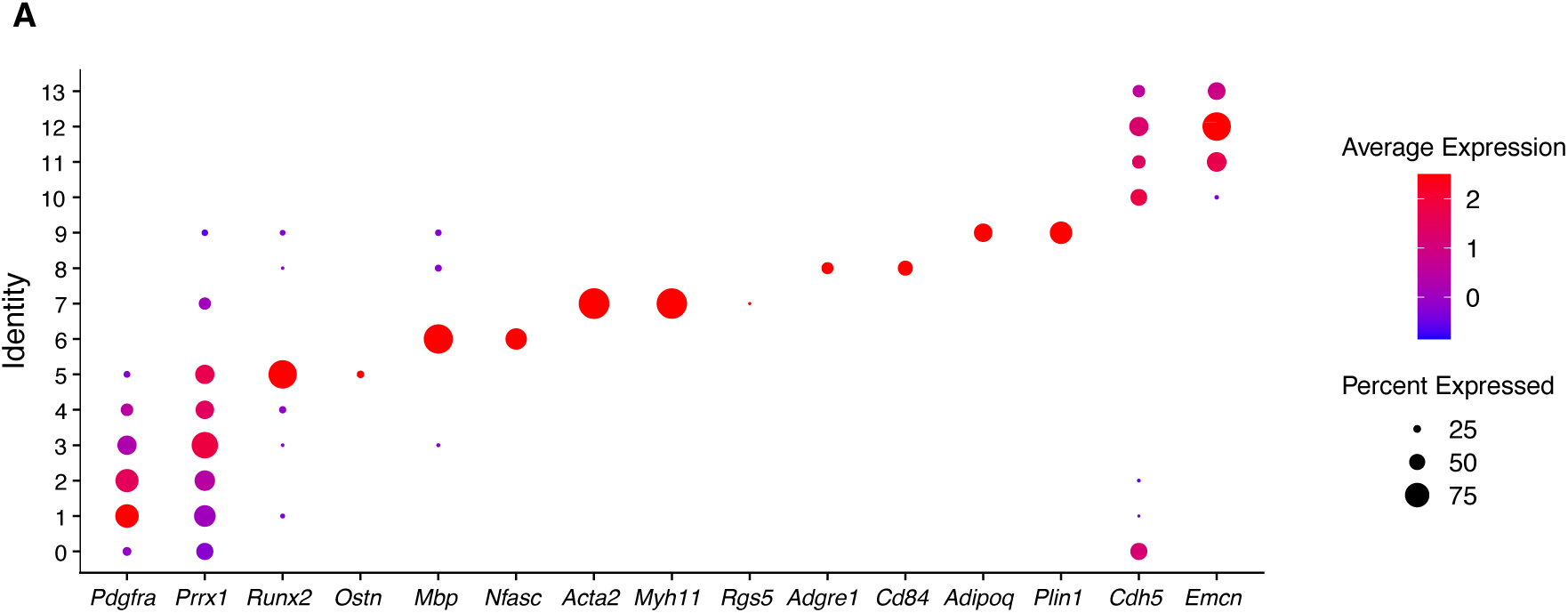
Dot plot of marker genes of the populations from uninjured periosteum.

**Figure 2 – Supplementary Figure 1:**
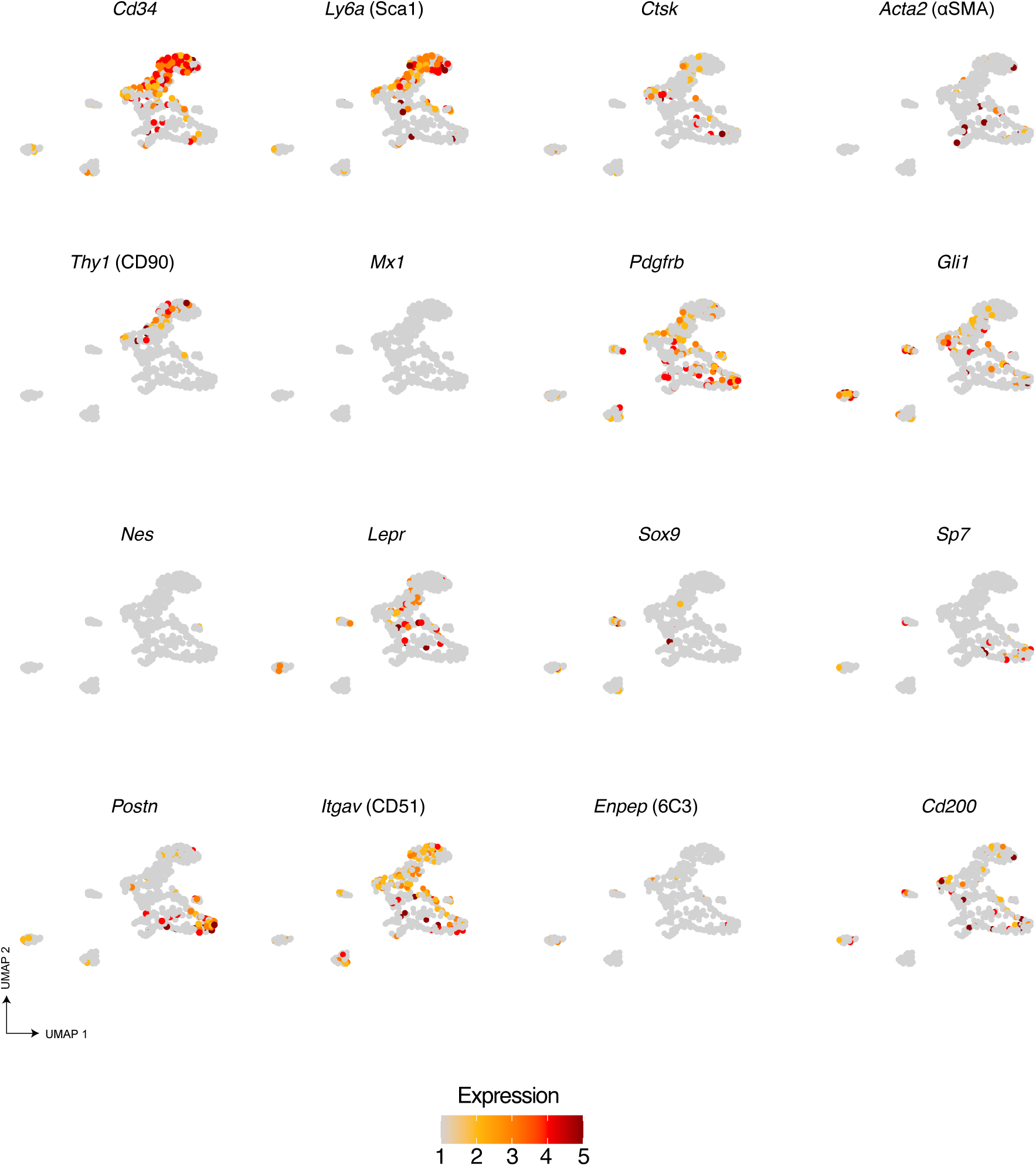
Expression of known SSPC markers in the periosteum at steady state. Feature plots of known markers of SSPCs in the SSPC/fibroblast subset in periosteum at steady state.

**Figure 3 – Supplementary Figure 1 :**
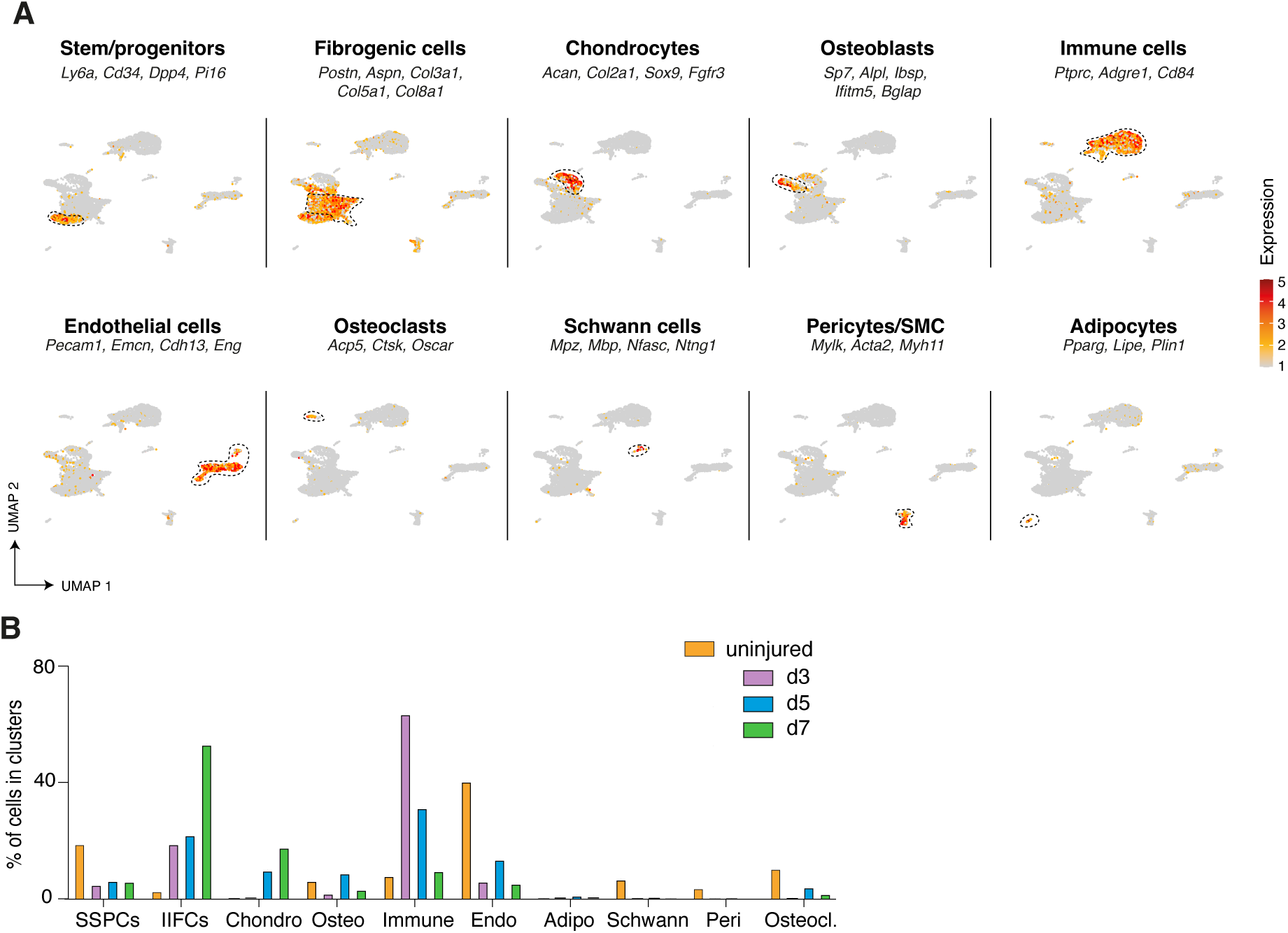
Heterogeneity and dynamics of the cell populations in the fracture environment. **A.** Feature plots of the lineage score of the different cell populations in the combined fracture datasets. **B.** Percentage of cells in each cell population per time point.

**Figure 3 – Supplementary Figure 2:**
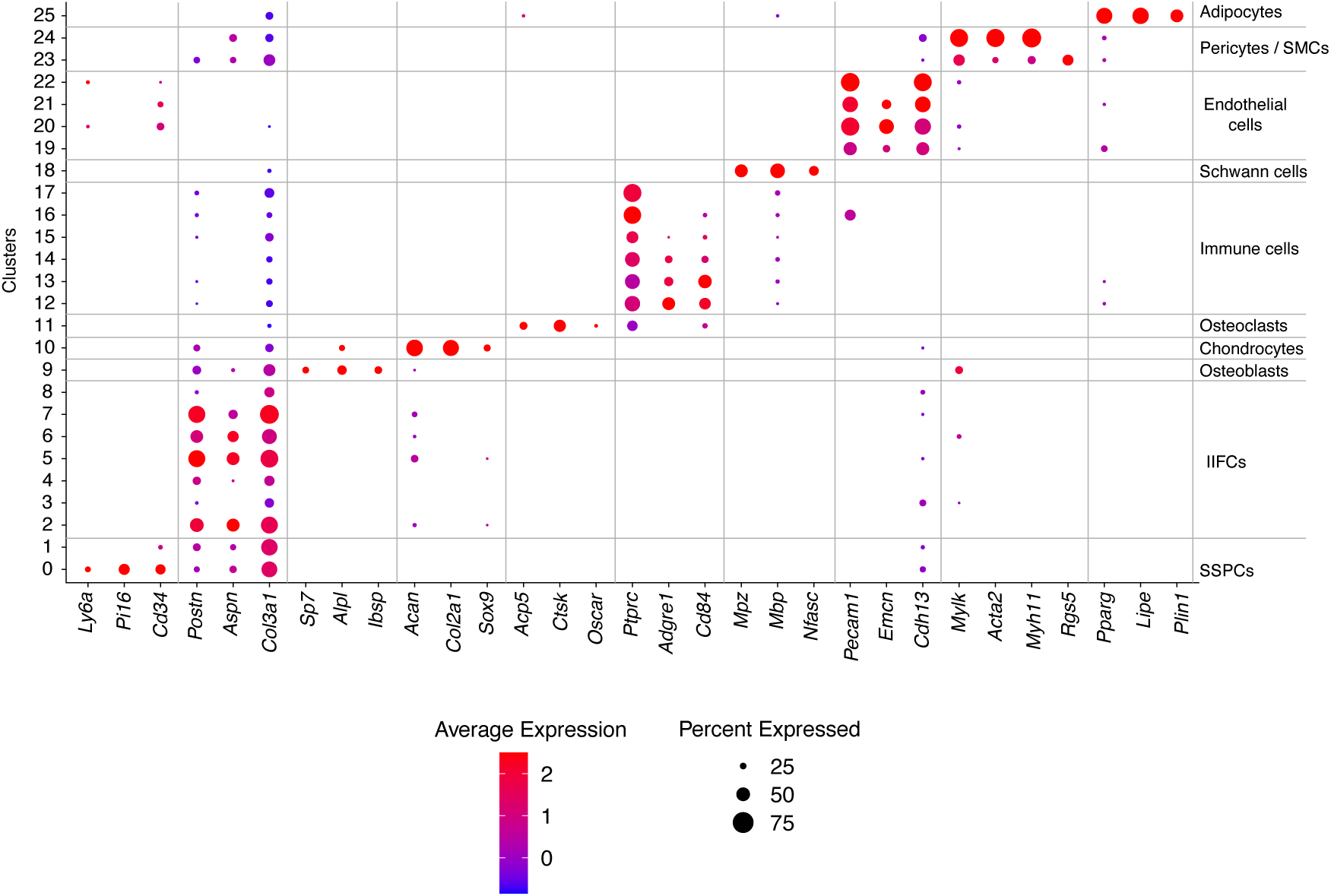
Dot plot of marker genes of the populations from the combined fracture dataset.

**Figure 4 – Supplementary Figure 1:**
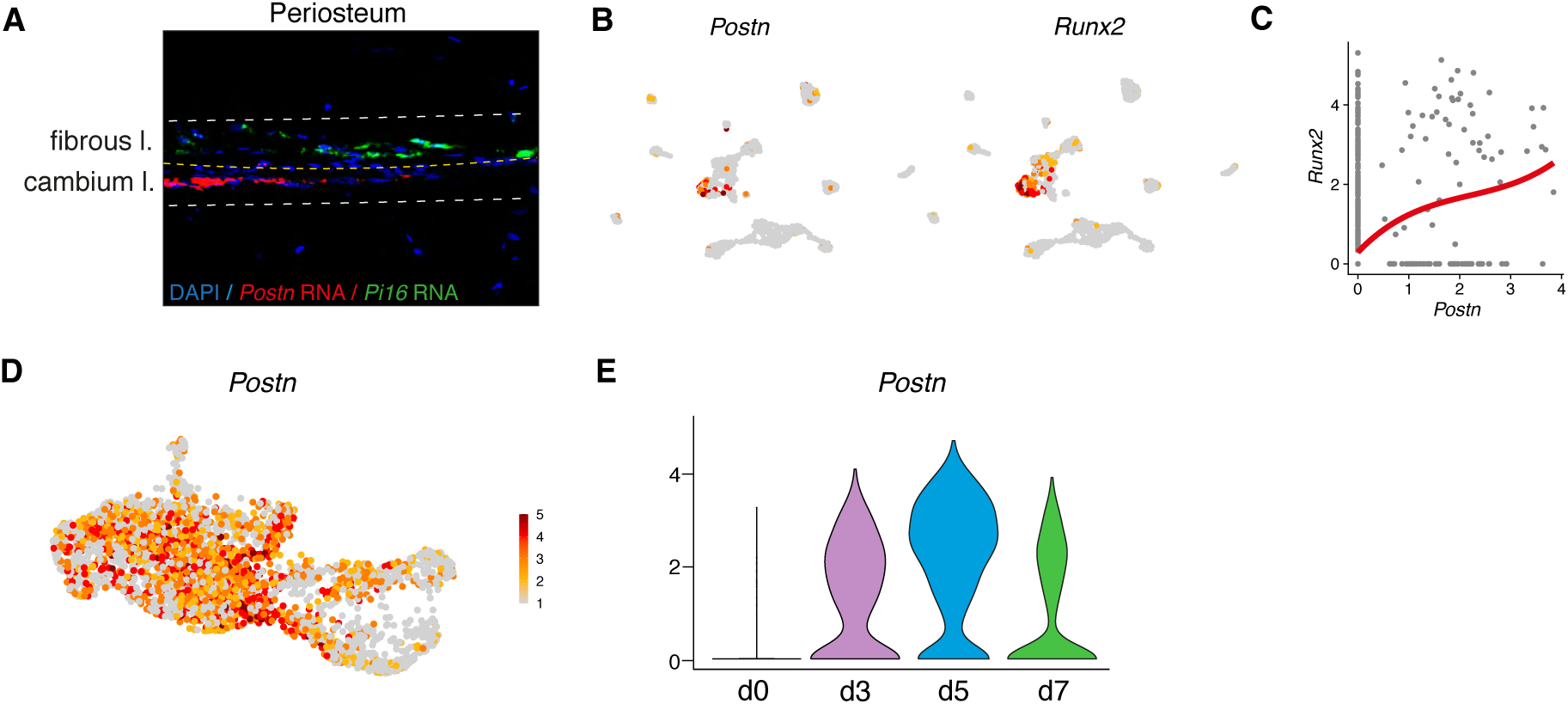
Periosteal SSPCs do not express Postn. **A.** RNAscope experiment showing the presence of *Postn* expressing cells in the inner cambium layer (cl) of the periosteum and *Pi16*-expressing cells in the fibrous layer (fl). **B.** Feature plots of *Postn* and *Runx2* expression in the uninjured periosteum. **B.** Scatter plot of *Runx2* and *Postn* expression in the uninjured periosteum dataset showing that *Postn* is mostly expressed by cells expressing *Runx2*. **D.** Feature plot of *Postn* expression in the subset of SSPCs, IIFCs, osteogenic and chondrogenic cells from Fig. 5B. **E.** Violin plot of *Postn* expression per time point.

**Figure 4 – Supplementary Figure 2:**
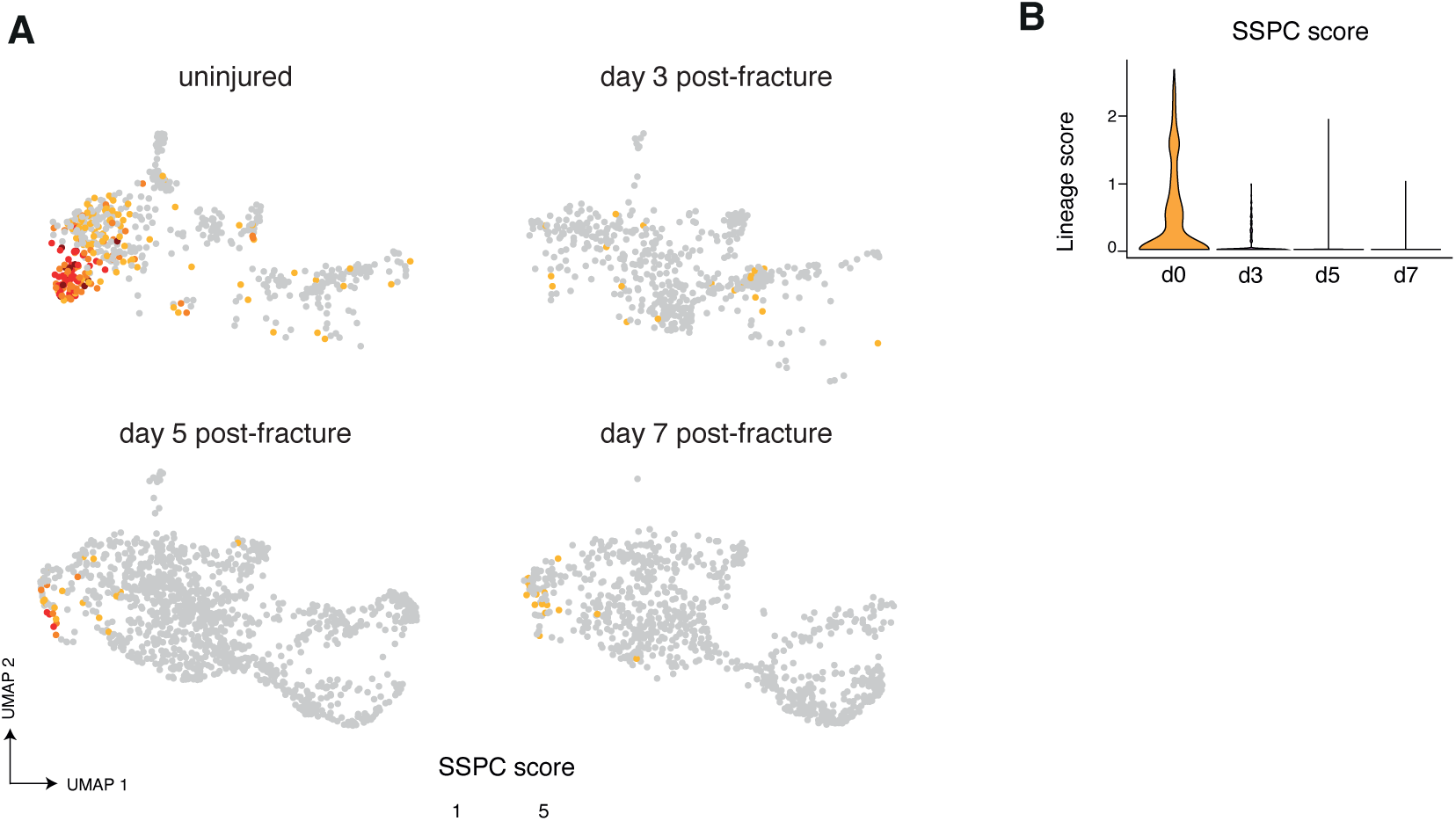
Absence of SSPCs in the injured periosteum. **A**. Feature plot of SSPC lineage score in the subset of SSPCs, IIFCs, osteoblasts and chondrocytes separated by time point from Fig. 5B. **B**. Violin plot of SSPC lineage score by time point.

**Figure 5 – Supplementary Figure 1:**
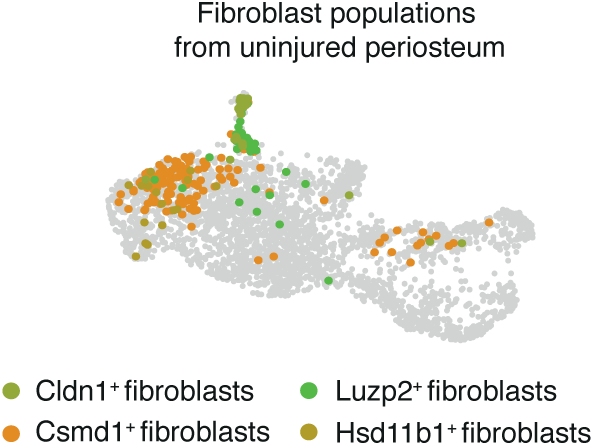
UMAP projection highlighting the distribution of periosteal fibroblasts in the combined fracture dataset.

**Figure 6 – Supplementary Figure 1:**
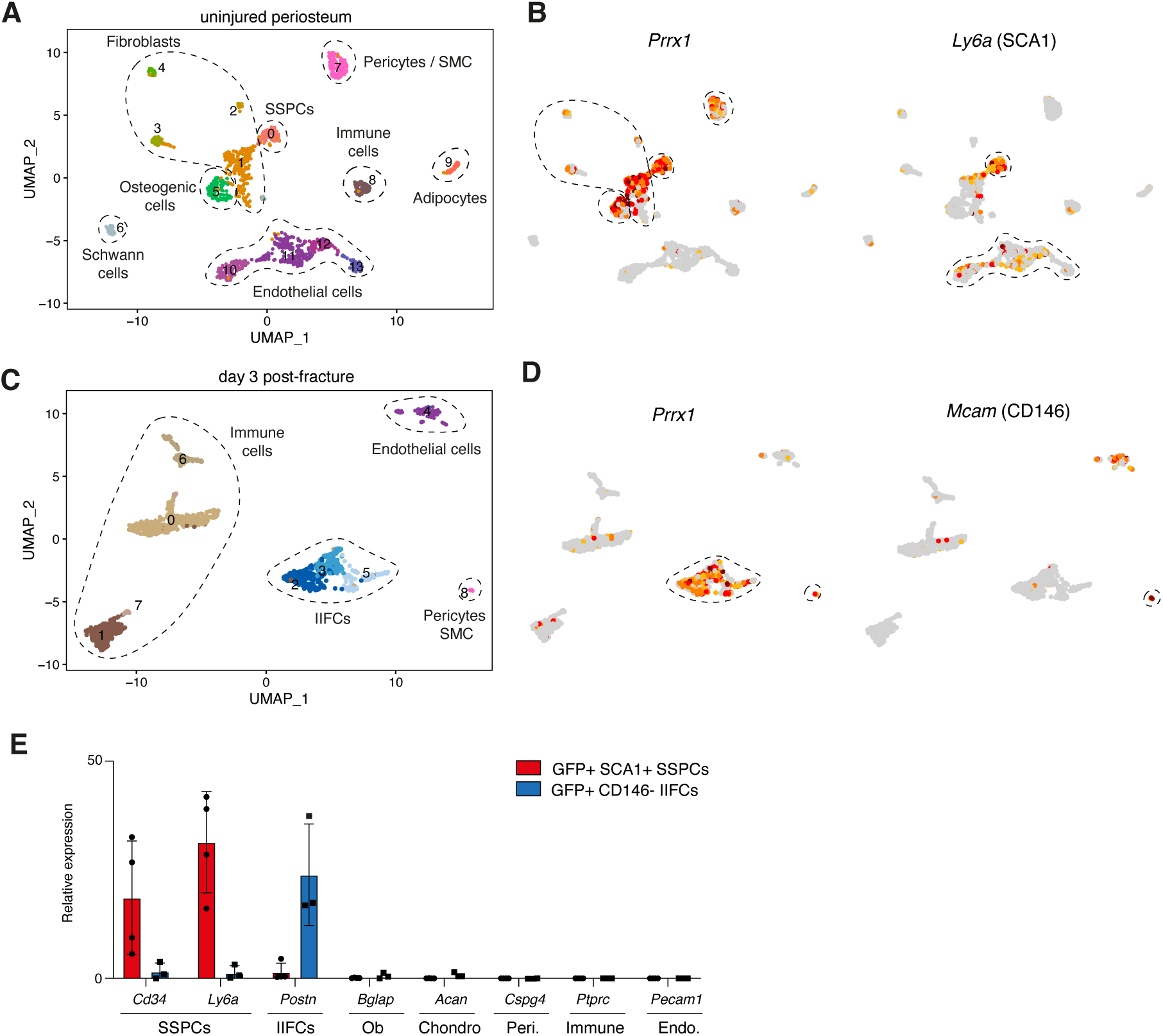
Validation of SSPC and IIFC sorting strategies. **A**. UMAP projection of the clustering of the uninjured periosteum. **B**. Feature plots of Prxx1 and Ly6a expression in the uninjured dataset. **C**. UMAP projection of the clustering of the day 3 post-fracture periosteum and hematoma. **D.** Feature plots of Prxx1 and Mcam (CD146) expression in the day 3 post-fracture periosteum and hematoma dataset. **E**. Relative expression of cell population markers by Prx1^+^ SCA1^+^ SSPCs from uninjured periosteum and Prx1^+^ CD146^-^ IIFCs from day 3 post-fracture periosteum and hematoma.

**Figure 7 – Supplementary Figure 1:**
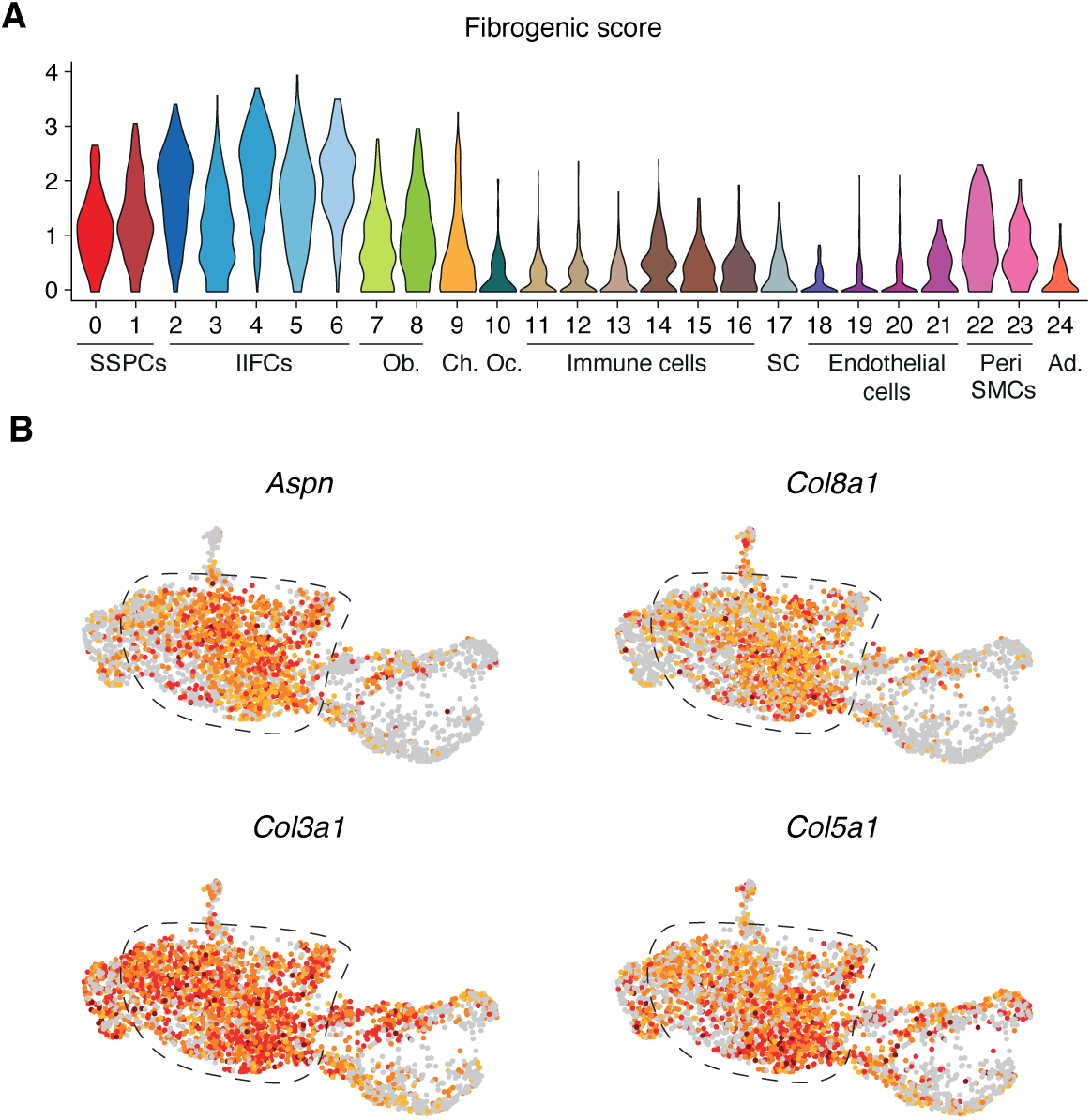
IIFCs are expressing ECM-related genes. **A.** Violin plot of the extracellular matrix genes score in the integrated dataset. **B**. Feature plots of *Aspn, Col3a1, Col5a1* and *Col8a1* in the subset of SSPCs, IIFCs, osteoblasts and chondrocytes.

**Figure 7 – Supplementary Figure 2:**
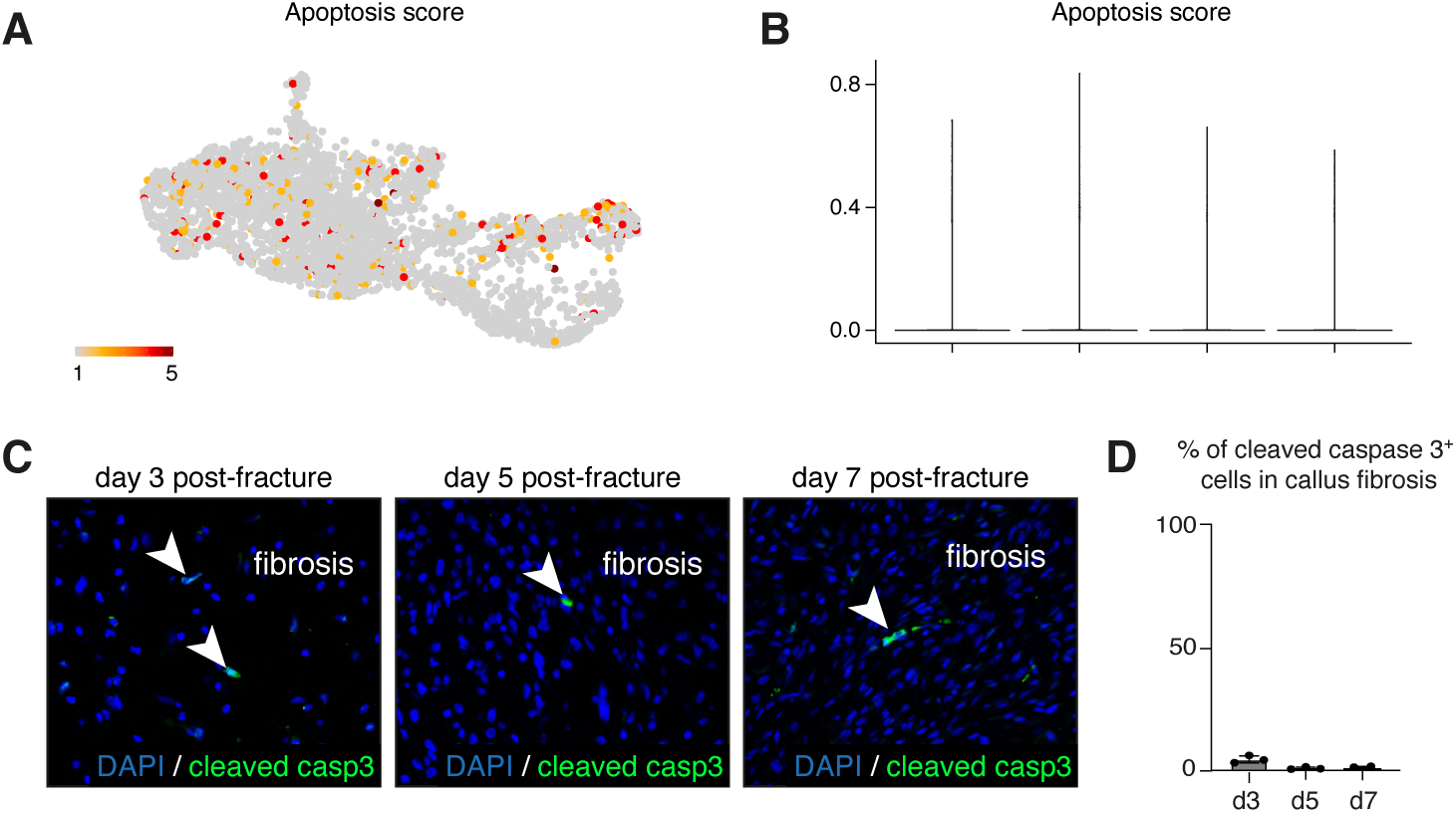
IIFCs do not undergo apoptosis. **A**. Feature plot of the apoptosis score in the subset of SSPCs, IIFCs, osteoblast and chondrocytes. **B**. Violin plot of the apoptosis score separated by time points. **C**. Immunofluorescence of cleaved caspase 3 in callus fibrosis at day 3, 5 and 7 post-fracture. **D**. Percentage of cleaved caspase 3 positive cells in the fibrosis callus fibrosis at day 3, 5 and 7 post-fracture.

**Figure 8 – Supplementary Figure 1:**
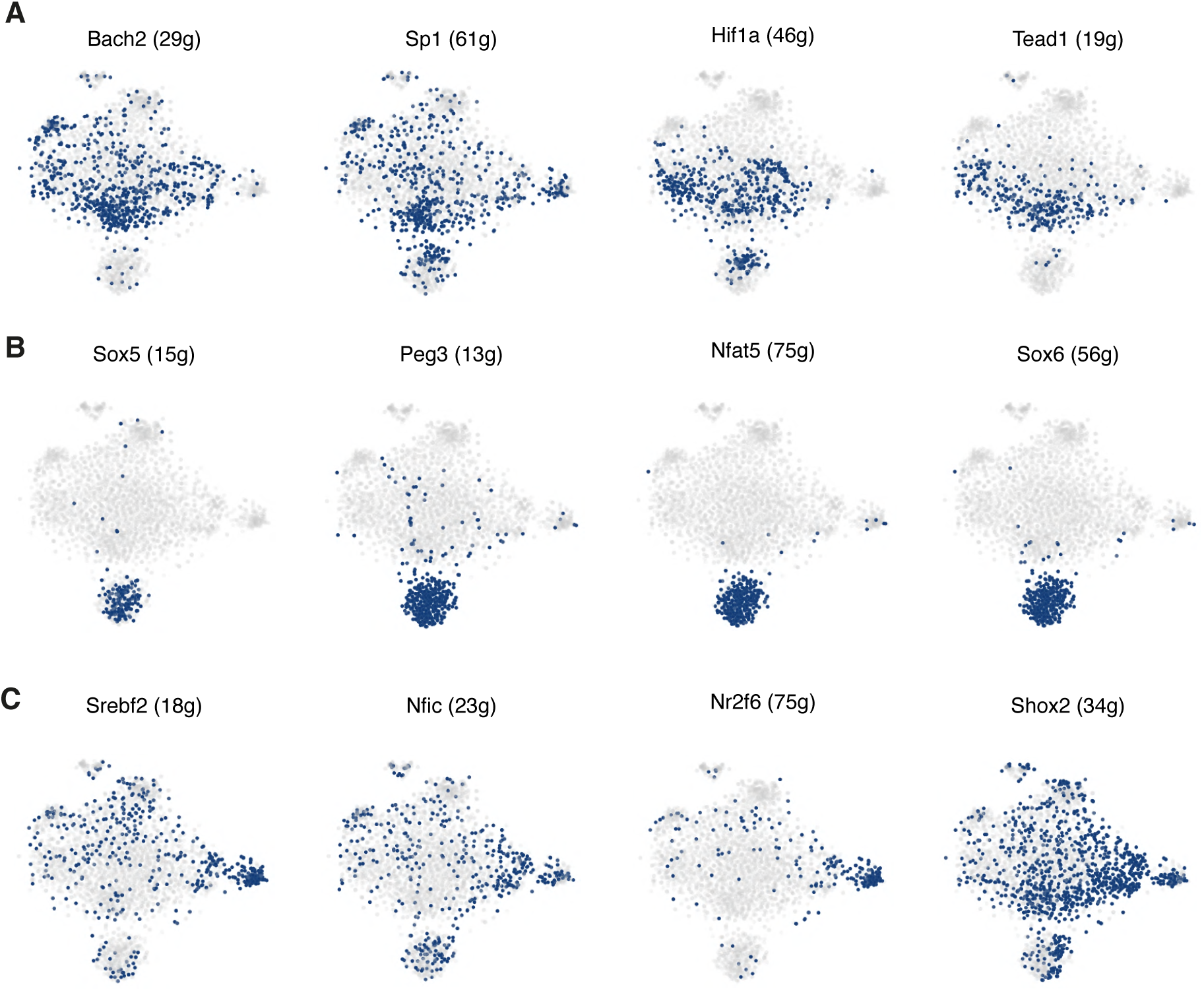
Regulon activity in the subset of SSPCs, IIFCs, osteoblasts and chondrocytes. **A.** Activity of Mta3, Six1, Sox9 and Sp7 regulons in the UMAP Seurat clustering of SSPCs, IIFCs, chondrocytes and osteoblasts. **B.** Activity of chondro-core 1 regulons in SCENIC tSNE clustering of SSPCs, IIFCs, chondrocytes and osteoblasts. **C.** Activity of chondro-core 2 regulons in SCENIC tSNE clustering of SSPCs, IIFCs, chondrocytes and osteoblasts. **D.** Activity of osteo-core regulons in SCENIC tSNE clustering of SSPCs, IIFCs, chondrocytes and osteoblasts.

**Figure 9 – Supplementary Figure 1:**
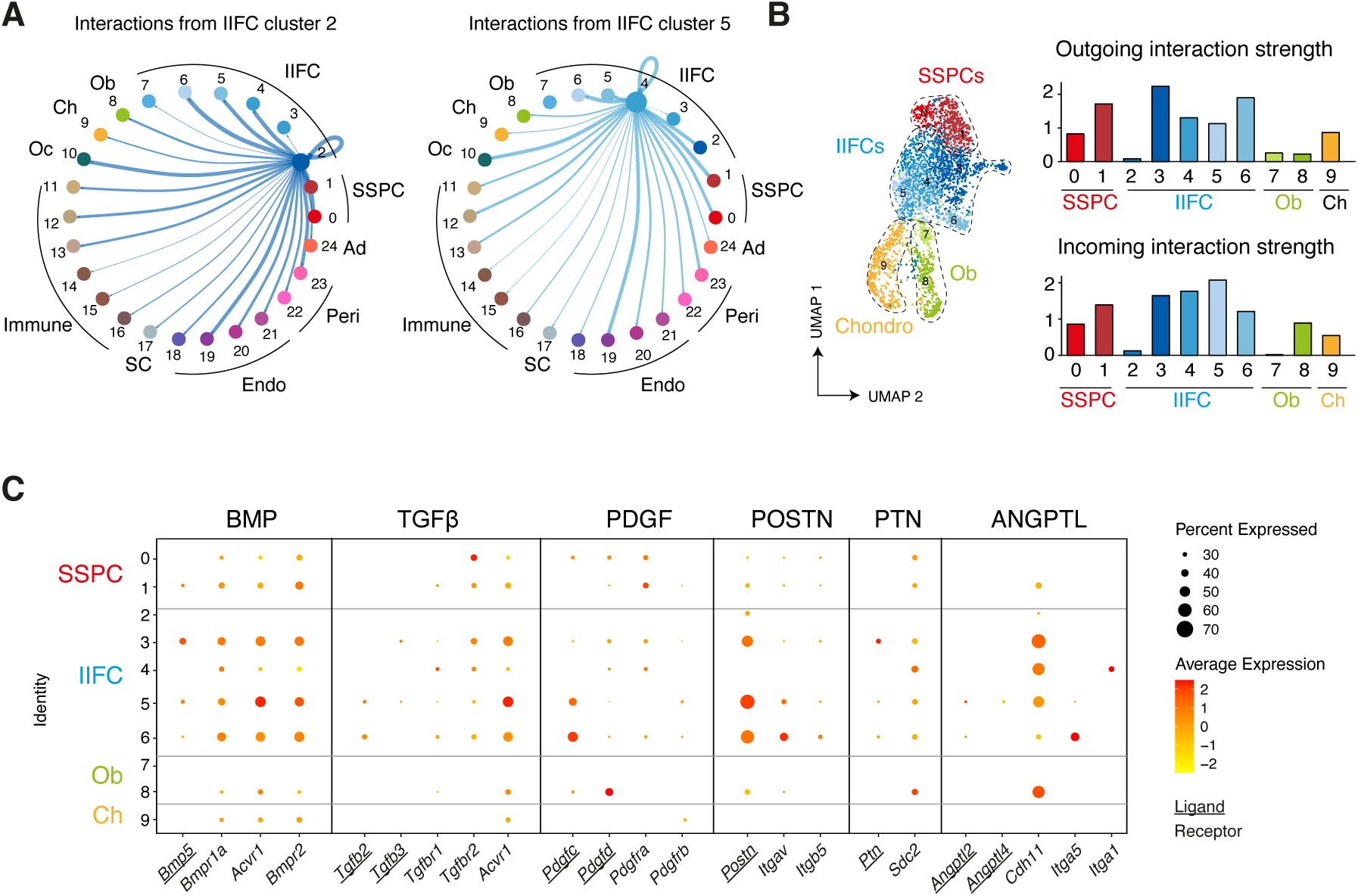
Paracrine interactions from IIFCs. **A.** Circle plots showing the interaction strengths between IIFCs in clusters 2 and 5 with the other cell populations. **B.** (left) Feature plot of the subset of SSPCs, IIFCs, chondrocytes and osteoblasts from Fig 5. (right) Outgoing and incoming interaction strengths of the subset of SSPCs, IIFCs, chondrocytes and osteoblasts. **C.** Dot plots of the expression of the ligands (underlined) and receptors of BMP, TGFβ, PDGF, POSTN, PTN and ANGPTL family involved in cell-cell interactions from IIFCs after fracture.

**Table S1:** Lists of genes used for lineage score analyses of murine snRNAseq.

| **Score** | **Genes** |
| --- | --- |
| **Stem/progenitor score** | *Ly6a, Cd34, Dpp4, Pi16* |
| **Fibrogenic score** | *Postn, Aspn, Col3a1, Col5a1, Col8a1* |
| **Chondrogenic score** | *Acan, Col2a1, Sox9, Fgfr3* |
| **Osteogenic score** | *Sp7, Alpl, Ibsp, Ifitm5, Bglap* |
| **Notch score** | *Notch1, Notch3, Notch2, Notch4, Jag1, Dtx3, Fbxw7, Psen1, Epn2, Sel1l, Maml1, Spen, Anxa4* |
| **ECM score** | *Col5a1, Col5a2, Col5a3, Col3a1, Col8a1, Col8a2, Col12a1, Eln, Postn, Aspn, Lox, Tll1, Smoc1, Pxdn, Smoc2* |
| **BMP score** | *Bmp5, Bmpr1a, Acvr1, Bmpr2* |
| **TGFb score** | *Tgfb2, Tgfb3, Tgfbr1, Tgfbr2, Acvr1* |
| **PDGF score** | *Pdgfc, Pdgfd, Pdgfra, Pdgfrb* |
| **POSTN score** | *Postn, Itgav, Itgb5* |
| **PTN score** | *Ptn, Sdc2* |
| **ANGPLT score** | *Angptl2, Angptl4, Cdh11, Itga1, Itga5* |
| **Total ligand score** | *Bmp5, Tgfb2, Tgfb3, Postn, Ptn, Angptl4, Angptl2, Pdgfc, Pdgfd* |
| **Total receptor score** | *Bmpr1a, Acvr1, Bmpr2, Tgfbr1, Tgfbr2, Itgav, Itgb5, Sdc2, Ncl, Pdgfra, Pdgfrb* |

**Table S2:** Lists of the regulons composing the cores.

| Group of regulons | Regulons |
| --- | --- |
| Fibro-core regulons | Arnt, Bach1, Bcl3, Cux1, E2f3, Elk4, Etv6, Fosl2, Foxn3, Foxp1, Jun, Junb, Mdb4, Meis1, Myc, Nfib, Pbx1, Pbx3, Rocor1, Runx1, Six1 |
| Chondro-core 1 regulons | Npas2, Hif1a, Sp1, Arntl, Bach2, Nfkb1, Maf, Tead1, Nfatc2 |
| Chondro-core 2 regulons | Sox6, Sox5, Sox9, Mef2c, Nfat5, Mef2d, Ccnt2, Clock, Ppard, Irf2, Trp53, Peg3, Creb3l2, Ets1 |
| Osteo-core regulons | Nr2f6, Nfic, Shox2, Tbx2, Sp7, Bcl11b, Tcf7, Runx2 |

**Table S3:** Top 5 terms from Reactome analysis on fibro-core regulons.

| Reactome number | Term | Adjusted p-value |
| --- | --- | --- |
| R-HSA-157118 | Signaling By NOTCH | 2,52E-04 |
| R-HSA-1912408 | Pre-NOTCH Transcription And Translation | 0,00250164 |
| R-HSA-1912422 | Pre-NOTCH Expression And Processing | 0,00331873 |
| R-HSA-5619507 | Activation Of HOX Genes During Differentiation | 0,00394002 |
| R-HSA-9018519 | Estrogen-dependent Gene Expression | 0,00697148 |

